# *Candida albicans* enhances the progression of oral squamous cell cancrinoma *in vitro* and *in vivo*

**DOI:** 10.1101/2021.03.31.437836

**Authors:** M Vadovics, N Igaz, R Alföldi, D Rakk, É Veres, B Szücs, M Horváth, R Tóth, A Szücs, P Horváth, L Tiszlavicz, C Vágvölgyi, JD Nosanchuk, A Szekeres, M Kiricsi, LG Puskás, A Gácser

**Author notes:** **Corresponding author** Gácser A.

## Abstract

Oral squamous cell carcinoma (OSCC) is a serious health issue worldwide. OSCC is highly associated with oral candidiasis, although it is unclear whether the fungus promotes the genesis and progression of OSCC or cancer facilitates the growth of the fungus. Therefore, we investigated whether *Candida* could directly influence OSCC development and progression. Our *in vitro* results suggest that the presence of live *C. albicans*, but not *C. parapsilosis*, enhances the progression of OSCC by stimulating the production of matrix metalloproteinases, oncometabolites, pro-tumor signaling routes, and overexpression of prognostic marker genes associated with metastatic events. We also found that oral candidiasis triggered by *C. albicans* enhanced the progression of OSCC *in vivo* through the induction of inflammation and overexpression of metastatic genes and markers of epithelial-mesenchymal transition. Taken together, these results suggest that *C. albicans* actively participates in the complex process of OSCC progression.

## Introduction

Oral cancer accounts for 2-4% of all cancer cases and includes a group of neoplasms that affects various regions of the oral cavity, the pharyngeal area, and salivary glands. It is estimated that more than 90% of all oral neoplasms are squamous cell carcinoma (OSCC)^1^. OSCC is the sixth most common cancer in the USA^2^ and has the highest mortality rate among all oral diseases and the lowest survival rates (nearly 50%) among all cancer types. It mainly occurs in individuals over 40^1^. Regarding worldwide prevalence, a 2012 study reported that oral cancer has the highest incidence in Hungary, Pakistan and India (7 or more cases per 100,000 individuals)^3^. Risk factors of oral cancer include poor oral hygiene, tobacco, alcohol, and meat consumption^4^. OSCC and other oral tumors are typically treated by surgery, radiation, and chemotherapy. Chemotherapy and radiotherapy, when used simultaneously, provide a synergistic benefit against OSCC^5^. Currently, chemoradiotherapy is the primary treatment for OSCC leading to adverse effects such as mucositis and myelosuppression^6^, which also affects the composition, quantity, and complexity of the oral microbiota^7–9^.

Being the most prevalent yeasts in the oral cavity^10–12^, *Candida* species, but especially *C. albicans*, are able to excessively proliferate and invade host mucosal tissues upon the disfunction or disruption of the epithelial barrier. Tissue invasion by *C. albicans* is due to the production and secretion of fungal hydrolytic enzymes, hypha formation, and contact sensing. While these phenotypic characteristics may endow *Candida* with a competitive advantage in the oral cavity, it is the host’s immune competence that ultimately determines whether clearance, colonization, or disease occurs^13^.

There is an association between the dysbiosis of oral yeasts and OSCC. Previous studies have revealed a higher yeast carriage and diversity in oral cancer patients compared to healthy individuals, and fungal colonisation in the oral cavity bearing OSCC is higher on the neoplastic epithelial surface compared to adjacent healthy surfaces indicating a positive association between oral yeast carriage and epithelial carcinoma^14–18^. These findings and other publications suggest that *Candida* cells may have a direct contribution to oral tumor development. One example is a case study demonstrating a rare event where persistent oral candidiasis led to the development of OSCC in an elderly patient^19^. Several other studies have confirmed the hyperplastic response of the epithelium when invaded by *Candida*. If the lesions caused by *Candida* are untreated, a minor proportion of epithelial cells may undergo dysplasia and transform into carcinoma. Thus, there is strong evidence supporting the idea that *Candida* contributes to carcinogenesis events in the oral cavity^20–24^. Additionally, patients undergoing radiotherapy for head and neck cancer are at increased risk of developing oral candidiasis as fungal colonization occurs more frequently in these patients ^25,26^. One study found that treatment of OSCC by chemoradiotherapy resulted in oral candidiasis in 75.3% of patients^27^. Several other studies demonstrated a link between OSCC and *Candida* colonisation of the mouth^23,28–31^. *Candida* infection in patients with cancer is usually considered to be the consequence of an altered immune status because both myelosuppression and mucositis enable the development of oral candidiasis^7,8,32,33^.

To date, there is increasingly strong evidence that suggests that the development of oral candidiasis in oral tumor patients enhances progression events that could result in poor prognosis. Despite of this, no study has effectively investigated this phenomenon and characterized the potential underlying mechanisms for *Candida* enhancing OSCC development and progression. In this study, we aimed to investigate fungal-specific molecular mechanisms that could facilitate and enhance cancer development or progression.

## Results

### Heat-inactivated *Candida* and zymosan slightly increase activity related to metastasis *in vitro*

To examine the effect of the increased fungal burden on oral tumor progression, OSCC cells HSC-2 and HO-1-N-1 were treated either with zymosan, heat-inactivated (HI) *C. albicans* or HI-*C. parapsilosis* cells. We used zymosan -a cell wall component of *Saccharomyces cerevisiae* – as a general fungal stimulus and HI-*Candida* treatment on the OSCC cells for examining the effects of direct contacts between fungal cell wall components and OSCC cells. First, a Wound Healing Assay was used to analyze the invasive capacity of OSCC cells. The cellular movement of the HO-1-N-1 cell line was significantly enhanced by all applied treatments compared to the untreated control (zymosan:1.445+/-0.076; HI-*C. albicans*:1.369+/-0.09; HI-*C. parapsilosis*: 1.454+/-0.083). In contrast, no significant differences were observed in the case of HSC-2 cells (Figure 1A). To assess host cell proliferation during fungal exposure, we performed BrdU proliferation assays. The BrdU assay revealed that HI-*Candida* cells do not affect tumor cell proliferation of either OSCC cell lines (Supplementary 1A).

**Figure 1.**
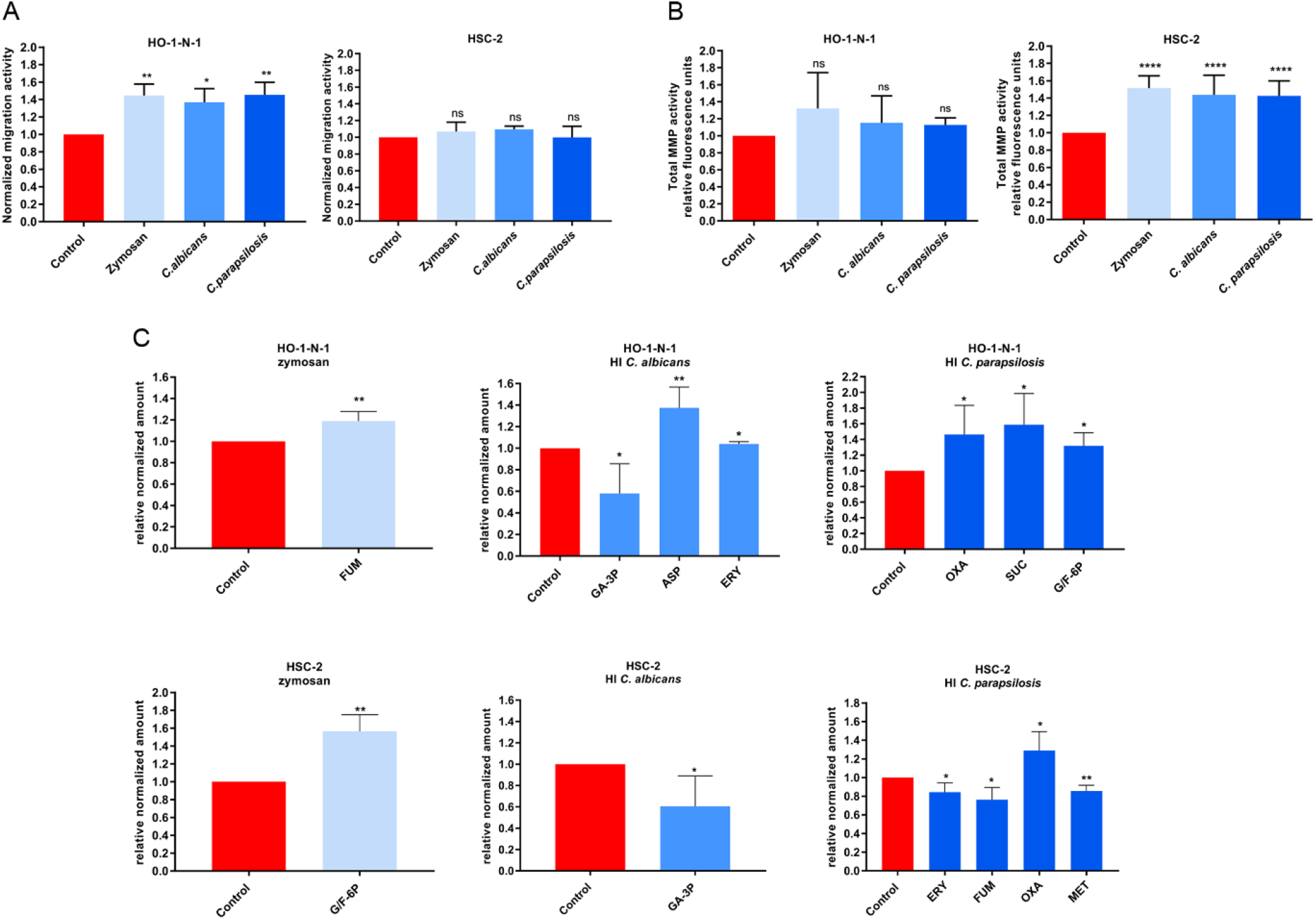
Effects of HI-*Candida* and zymosan on HO-1-N-1 and HSC-2 oral squamous cell carcinoma cells *in vitro*. (**A**) Normalized migration activity of OSCC cells in the presence of HI-*C. albicans*, HI-*C parapsilosis* and zymosan measured by a Wound Healing Assay, n=3. (**B)** Normalized total secreted matrix metalloproteinase (MMPs) activity of OSCC cells in the presence of HI-*C. albicans*, HI-*C. parapsilosis* and zymosan measured by a total MMP activity kit, n=4. (**C**) Normalized amounts of metabolites of OSCC cells in the presence of HI-*C. albicans*, HI-*C. parapsilosis* and zymosan as measured by HPLC-HRMS, n=4; FUM – Fumaric acid; GA-3P -Glyceraldehyde-3P; ASP -Aspartic acid; ERY -Erythrose-4P, OXA -Oxaloacetic acid; SUC -Succinic acid; G/F-6P – Glucose/Fructose-6p; MET –Methionine; Control: tumor cells without any treatment. Unpaired t-test; * p ≤ 0.05; ** p ≤ 0.01; *** p ≤ 0.001; **** p ≤ 0.0001

A critical component of tumor cell adaptability to different environment lies in their ability to remodel the extracellular matrix of the surrounding tissue, which is largely attributed to the secretion of a variety of proteases (serine, cysteine, threonine, aspartic, and metalloproteinases). In particular, matrix metalloproteinases (MMPs) are key enhancers of tumor dissemination^34^. To assess whether yeast presence affects MMP secretion and function, secreted total MMP activity of the OSCC cell lines was measured. In all applied conditions significantly elevated secreted MMP activity was observed in the case of HSC-2 cells compared to the untreated control (zymosan: 1.516+/-0.041; HI-*C. albicans*: 1.437+/-0.06536; HI-*C. parapsilosis*: 1.426+/-0.057), however, the MMP activity was not altered significantly when HO-1-N-1 cells were exposed to treatments (Figure 1B).

Metabolites generated by cancer cells influence the metastatic cascade, affecting epithelial-mesenchymal transition (EMT), the survival of cancer cells in circulation, and metastatic colonization at distant sites^35^. Potential changes in metabolic activity were examined by analyzing the amount of glycolysis and TCA cycle intermediates, and of certain amino acids (Supplementary 1C) with HPLC-HRMS. In the case of HO-1-N-1 cells, zymosan treatment produced a significant change in fumaric acid (1.189+/-0.044), while HI-*C. albicans* treatment altered the levels of glyceraldehyde-3P (0.580+/-0.137), aspartic acid (1.374+/-0.096), and erythrose-4P (1.039+/-0.015). HI-*C. parapsilosis* treatment altered the production of oxaloacetic acid (1.463+/-0.214), succinic acid (1.586+/-0.199), and glucose/fructose-6p (1.317+/-0.118) (Figure 1C). For the HSC-2 cell line, zymosan treatment significantly increased the production of glucose/fructose-6p (1.564+/-0.132), while HI-*C. albicans* treatment reduced the concentrations of glyceraldehyde-3P (0.605+/-0.142). HI-*C. parapsilosis* treatment altered levels of erythrose-4P (0.844+/-0.056), fumaric acid (0.763+/-0.065), oxaloacetic acid (1.289+/-0.117), and methionine (0.855+/-0.031) (Figure 1C). Non-significant changes are shown in supplementary figure 1C.

Taken together, HI-*Candida* and zymosan treatment had a significant effect on migration, secreted MMP activity, and oncometabolite production of OSCC cells, which suggests interactions between the tumor cells and the components of fungal pathogen-associated molecular patterns (PAMPs).

### Live *Candida* enhances detachment, MMP activity, and metabolite production of OSCC cells *in vitro*

After assessing the direct interactions, we aimed to examine the indirect interplay between OSCC cells and *Candida*. To accomplish this, we applied live *C. albicans* and *C. parapsilosis* cells on OSCC cultures and recorded the movement of cancer cells on time-lapse video. In the presence of live *C. albicans*, increased numbers of detached, single HSC-2 cells were detected around the fungal cells compared to untreated control cancer cells (Figure 2A). Live *C. parapsilosis* did not cause noticeable changes in the movement of HSC-2 cells (Supplementary video 1,2,3). No change was detected in any of the fungal treatments in case of HO-1-N-1 cell line (data not shown). Similar to HI-*Candida*, live *C. albicans* and *C. parapsilosis* did not cause any changes in the proliferation of the OSCC cells as measured by BrdU assay (Supplementary 1B). Secreted total MMP activity was increased in both cancer cell lines following incubation with live *C. albicans* (HSC-2: 1.918+/-0.209; HO-1-N-1: 1.918+/-0,183), but the MMP activity in the presence of live *C. parapsilosis* was similar to that of the untreated control (Figure 2B). Metabolic changes of OSCC cells were also measured following treatments with live *C. albicans* and *C. parapsilosis*. No fungal metabolites could be measured after our extraction method. In the case of the HSC-2 cell line, live *C. albicans* treatment led to significant changes in the secretion of aspartic acid (2.67+/-0.346), glyceraldehyde-3P (0068+/-0.009), methionine (0,634+/-0.063), proline (0.666+/-0.087), and succinic acid (1.975+/-0.234), while *C. parapsilosis* treatment resulted in alterations in fumaric acid (1.199+/-0.046), glyceraldehyde-3P (0.225+/-0.076), and succinic acid (1.523+/-0.101) secretion. For the HO-1-N-1 cell line, *C. albicans* treatment significantly altered the secretion of aspartic acid (5.526+/-1.667), glyceraldehyde-3P (0.543+/-0.038), glucose/fructose-6p (0.288+/-0.047), glutamine (2.616+/-0.667), α-ketoglutaric acid (1.532+/-0.051), and succinic acid (9.81 +/-2.709), while *C. parapsilosis* treatment reduced the level of aspartic acid (0.408+/-0.024) (Figure 2C). The results demonstrated that live *Candida* caused prominent changes in the movement, MMP activity, and metabolite production in OSCC cells compared to HI-*Candida* and zymosan treatment.

**Figure 2.**
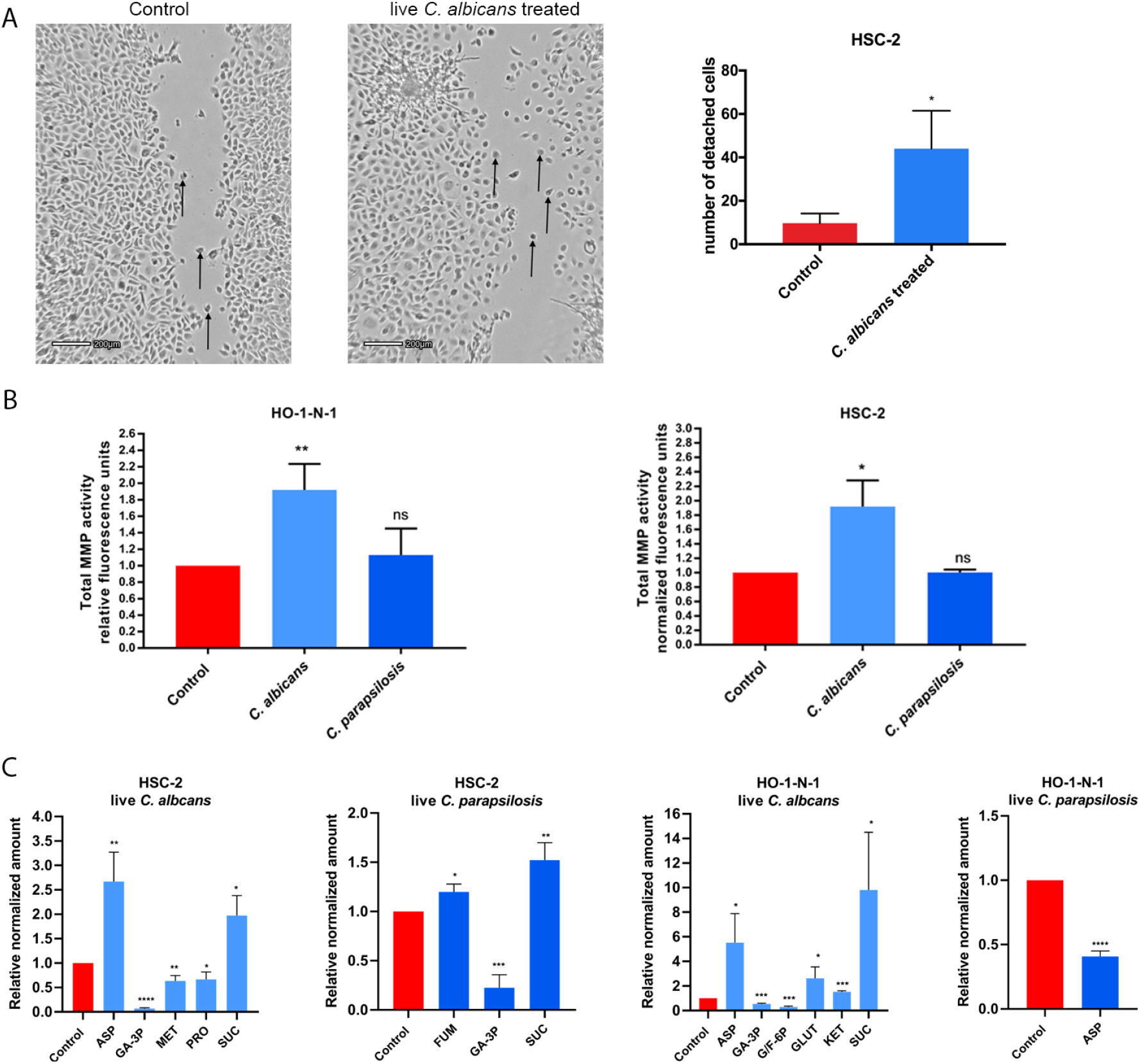
Effects of live *Candida* on HO-1-N-1 and HSC-2 oral squamous cell carcinoma cells *in vitro*. (**A**) Pictures from time-lapse videos of cellular migration of HSC-2 cells, arrows are pointing to detached cancer cells. The left picture shows the control cells and the right shows the live *C. albicans* treated cells. The graph shows the number of detached cells, n=3. (**B**) Normalized total secreted matrix metalloproteinase activity of OSCC cells in the presence of live *C. albicans* and *C. parapsilosis* as obtained by a total MMP activity kit, n=3. (**C**) Normalized amounts of metabolites of OSCC cells in the presence of live *C. albicans* and live *C. parapsilosis* as measured by HPLC-HRMS, n=3. ASP -Aspartic acid; GA-3P -Glyceraldehyde-3P; MET –Methionine; PRO – Proline; SUC -Succinic acid; FUM -Fumaric acid; G/F-6P – Glucose/Fructose-6p; GLUT -Glutamic acid; KET -α-Ketoglutaric acid. Control: tumor cells without any treatment. Unpaired t-test; * p ≤ 0.05; ** p ≤ 0.01; *** p ≤ 0.001; **** p ≤ 0.0001

### Live *C. albicans* stimulus activates genes and signaling pathways involved in the metastatic processes of OSCC

To identify and compare the molecular mechanisms of the observed cellular effects caused by each treatment, whole transcriptome analysis was performed with HSC-2 and HO-1-N-1 cells following exposure to zymosan, HI-*C. albicans*, HI-*C. parapsilosis*, live *C. albicans*, and live *C. parapsilosis*. The transcriptome analyses revealed that the highest number of statistically significant expression changes in OSCC were induced by live *C. albicans* treaments (HSC-2: n=2764; HO-1-N-1: n=137), followed by zymosan stimulus (n=19), while live *C. parapsilosis* and heat-inactivated fungal challenge did not trigger a significant response on the gene expression level in either of the OSCC cell lines. When comparing the OSCC cell lines, the evoked host response was more significant in HSC-2 (upregulated genes n=1315; downregulated genes n=1449) than in HO-1-N-1 cells (upregulated n=134; downregulated n=3) (Figure 3A) (Supplementary table2). Moreover, subsequent analyses suggest that *C. albicans* stimulus triggered significant changes in the expression of those HSC-2 genes (32 upregulated, 4 downregulated) that were previously described as marker genes of OSCC invasion (Supplementary table 4) (Figure 3B). Additional analyses provided another set of genes (21 upregulated, 3 downregulated), which overlapped with the characteristic profile of epithelial-mesenchymal transition derived from single-cell sequencing (scSeq) results from 18 patients with head and neck squamous cell carcinoma (HNSCC)^36^ (Figure 3C). We found 5 genes, INHBA, MMP10, MMP1, SEMA3C, and FHL2, which were present both in the overlapping subset with the OSCC invasion marker genes and the scSeq derived gene set (Figure G). The analyses also revealed that even though the HO-1-N-1 response was modest compared to HSC-2, 13 OSCC invasion marker genes (Figure 3D) and four EMT genes (Figure E) were upregulated after live *C. albicans* stimulus. Two of these genes, SERPINE1 and INHBA, overlapped with the OSCC invasion marker genes and EMT subsets (Figure H).

**Figure 3.**
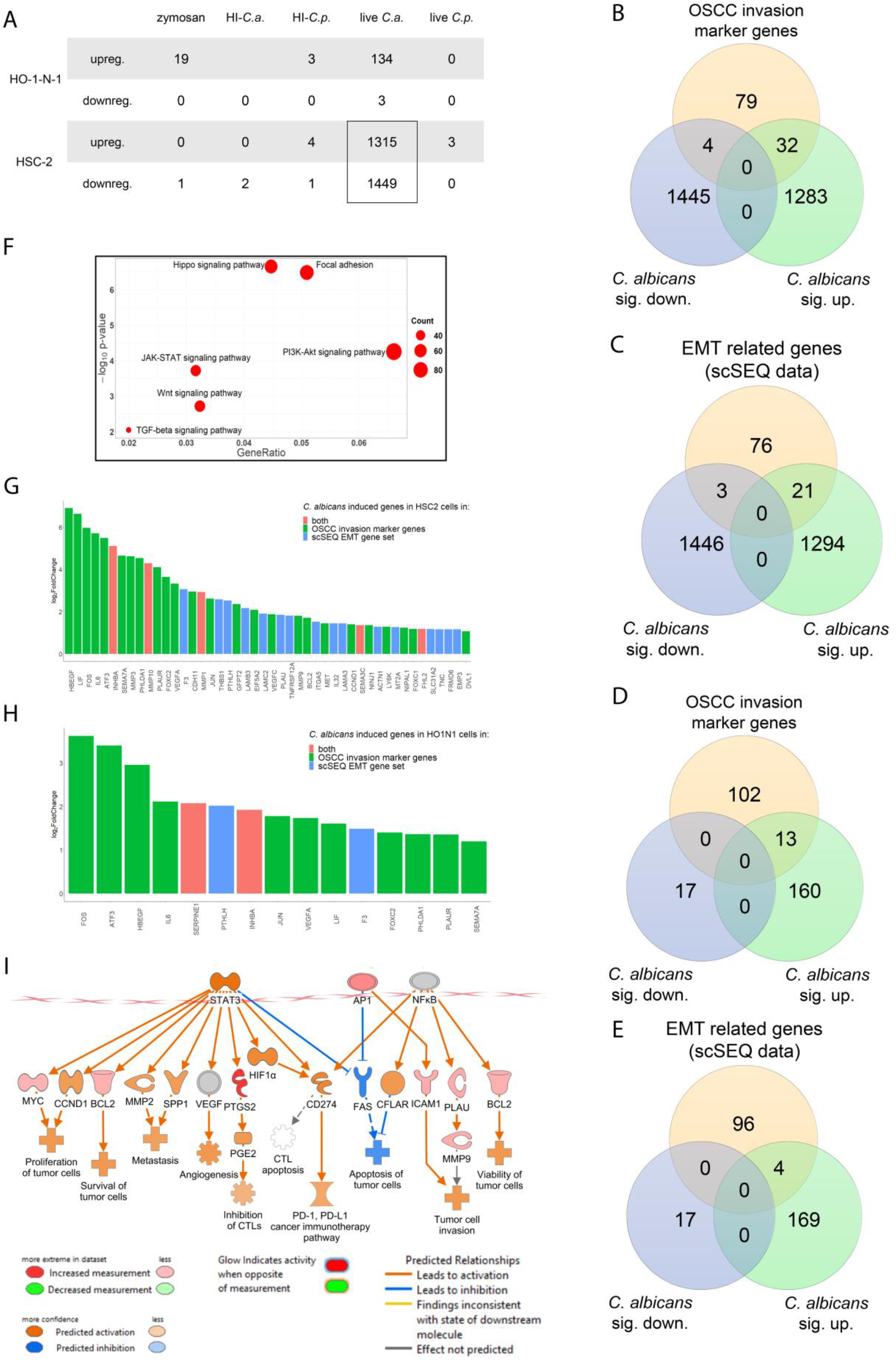
*In vitro* transcriptomic analysis. *Candida albicans* activates genes and signaling pathways involved in the OSCC metastatic processes. (**A**) Number of up or downregulated genes of HSC-2 and HO-1-N-1 cells after different fungal treatments, n=3. (**B**) VEN diagram of up or downregulated genes in HSC-2 cells in the presence of live *C. albicans* and OSCC invasion marker genes found in the literature. (**C**) VEN diagram of up or downregulated genes in the HSC-2 cell line in the presence of live *C. albicans* and EMT marker genes in HNSCC according to a single cell sequencing study. (**D**) VEN diagram of up or downregulated genes in HO-1-N-1 cells incubated with live *C. albicans* and OSCC invasion marker genes found in the literature. (**E**) VEN diagram of up or downregulated genes in the HO-1-N-1 cell line in the presence of live *C. albicans* and EMT marker genes in HNSCC according to a single cell sequencing study. (**F**) Signaling pathways that are key regulators of the OSCC invasion processes that were significantly activated in HSC-2 cells in the presence of live *C. albicans*. (**G**) Graph shows log2 fold change of *C. albicans* induced genes in HSC-2 cells involved in OSCC invasion according to the literature and of single-cell sequencing (scSeq) results from 18 patients with HNSCC. Red columns show OSCC marker genes according to the literature and scSEQ data. (**H**) Graph shows log2 fold change of *C. albicans* induced genes in HO-1-N-1 cells involved in OSCC invasion according to the literature and scSEQ data. Red columns show OSCC marker genes according to the literature and scSEQ data. (**I**) Causal analyses of the genes for which expression changed in HSC-2 cells after live *Candida albicans* treatment.

KEGG pathway analysis was also performed on the data set. It revealed significant activation of several pathways in HSC-2 cells that have previously been associated with OSCC metastasis development including the Hippo signaling pathway, Focal adhesion pathway, JAK-STAT pathway, PI3K-Akt pathway, Wnt pathway, and TGFβ pathway (Figure 3F, Supplementary 2-7)^37–41^. We analyzed further the expression data in the Ingenuity pathway analysis (IPA), Qiagen’s cutting-edge bioinformatical software. We employed the built-in causal analyses to investigate activation patterns of several intracellular signaling pathways based on the coherent regulation of their molecular elements. IPA analyses of the HSC-2 gene set predicted the activation of tumor-related pathways, including the tumor microenvironment pathway as well as the significant activation of several prognostic features, such as metastasis, invasion, angiogenesis, and proliferation of tumors based on the *C. albicans* stimulus-derived DEGs (Figure 3I).

Interestingly, the expression of DLST and SUCLA genes in the HSC-2 cell line was slightly increased in the presence of *C. albicans*, and these genes are involved in succinic acid metabolism. Furthermore, ASNSD1 and GOT1 genes were also slightly upregulated and they are involved in aspartic acid synthetic processes (Supplementary 9A). The identified genes and signaling pathway components upregulated in HSC-2 cells in the presence of live *C. albicans* were validated by qPCR (supplementary 7A-E).

These results support our previous findings that the presence of live *C. albicans* cells enhances the metastatic features of OSCC cells, which is in contrast to that found with live *C. parapsilosis*, HI-*Candida*, or zymosan.

### Establishment of a novel *in vivo* mouse xenograft model of OSCC and oral candidiasis

We successfully developed a novel *in vivo* mouse model of OSCC and oral candidiasis. In this model we aimed to thorougly examine the effect of oral candidiasis on OSCC progression. As *C. albicans* cells are not in direct contact with HSC-2 cells in this model, only an indirect effect of oral candidiasis can be examined on OSCC progression. To mimic the immunologic condition of patients caused by chemoradiotherapy, cortisone acetate was administered to 6-8 weeks old BALB/c mice to induce an immunosuppressed condition for subsequent tumor cell injection. As HSC-2 cells were more reactive to the presence of yeast cells compared to HO-1-N-1, 1×10^6^ cells of HSC-2 were injected in the tongues of mice to induce OSCC development. According to our experiments, a fully developed tumor formed on the tongues by day 3 after HSC-2 injection. After confirming the presence of a tumor, *C. albicans* saturated calcium alginate swabs (placed in to 1×10^9^/mL *C. albicans* suspension for 5 min) were placed under the tongue of each mouse for 75 minutes on day 5 (3 days after tumor cell injection). Mice were terminated on day 8 upon signs of decreased activity and weight loss (25%) to prevent unnecessary suffering (*Supplementary 10C*). On the 8^th^ day, the average tumor size was 5 mm in diameter (Figure 4B). Hematoxylin-eosin (H&E) staining was performed for histopathological analyses (Figure 4C). Fungal hyphae were detected on the mucosa following periodic acid–Schiff (PAS) staining (Figure 4D). The oral fungal burden of mice infected with *C. albicans* SC5314 was 10^5^–10^6^ CFU per g tissue, which is comparable with the literature of oral candidiasis models^42^.

**Figure 4.**
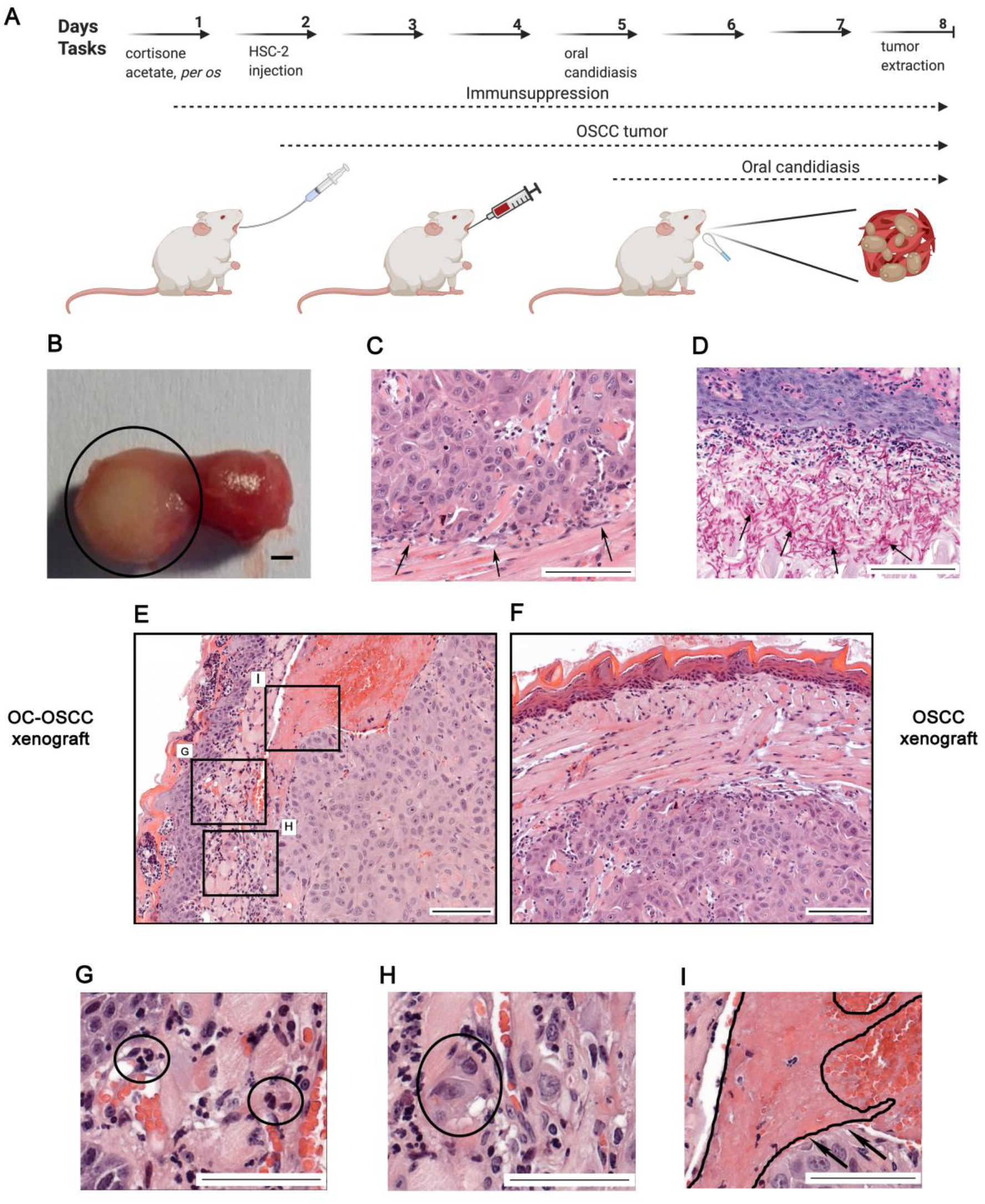
A new *in vivo* mouse model for the investigation of oropharyngeal candidiasis on the progression of OSCC. (**A**) Schematic figure of mouse xenograft for the investigation of the effect of *C. albicans* on the progression of OSCC. Immunosuppression and injection of human HSC-2 OSCC cells to the tongue of mice (OSCC xenograft). OSCC xenograft and oral candidiasis (OC-OSCC xenograft). The cartoon was produced by BioRender. (**B**) Representative mouse tongue on the 8^th^ day of the experiment (7 days after tumor cell injection). The circle highlights the tumor. Scale bar: 1 mm. (**C**) Histopathological image of the tumor on the 8^th^ day with arrows indicating the tumor edge. Scale bar: 100 µm. (**D**) Histopathological examination of the tumor on the 8^th^ day (7 days after tumor injection and 3 days post-infection) with black arrows indicating the fungal hyphae in the mucosa. Scale bar: 100 µm (**E**) Histopathological picture about the tumor on the 8^th^ day after HSC-2 injection and oral candidiasis (OC-OSCC xenograft, n=4 derived from four different experiments) Scale bar: 100 µm. (**F**) Histopathological picture about the tongue on the 8^th^ day after HSC-2 injection (OSCC xenograft, n=4 derived from four different experiments). Scale bar: 100 µm. (**G**) Infiltrating immune cells in OC-OSCC xenograft samples indicating *C. albicans* caused inflammation. Scale bar: 100 µm. (**H**) Detached budding tumor cells in OC-OSCC xenograft samples indicating epithelial to mesenchymal transition. Scale bar: 100 µm (**I**) Thrombosis in OC-OSCC xenograft samples. Scale bars: 100 µm.

### Oral candidiasis enhances the progression of OSCC *in vivo*

For investigating the effect of increased yeast burden on OSCC progression two animal groups were compared: one group received only cortisone acetate and HSC-2 human tumor cells (OSCC xenograft), while another group of animals received cortisone acetate, HSC-2 cells and *C. albicans* treatment for the development of oral candidiasis (OC-OSCC xenograft). Each group (OSCC and OC-OSCC xenograft) contained 4-4 animals derived from four different experiments. Histopathological samples of *Candida*-colonized and *Candida*-free tumors were analysed and scored manually in a blinded manner by a pathologist after H&E staining for the identification of inflammation, necrosis, infiltrating or pushing tumor edge, epithelial-mesenchymal transition (EMT), invasion markers, and signs and symptoms of thrombosis and peritumoral inflammation (Supplementary 10A). EMT and budding of tumor cells were more frequent in 2 samples out of 4 in the case of oral candidiasis compared to control tumors. Interestingly, thrombosis was also detected in 3 OC-OSCC xenograft samples. Only one sample showed thrombosis in OSCC xenograft.

Infiltrating immune cells were detected within the OC-OSCC xenograft samples, indicating *C. albicans*-induced inflammatory responses (*Figure 4G*). Images indicative of epithelial-mesenchymal transition were also identified, as represented by the increased number of detached individual tumor cells (Figure 4H). Interestingly, significant thrombosis was noted on the histopathological samples derived from OC-OSCC xenograft mice (*Figure 4I*).

Taken together, histopathological scoring suggests the presence of oral candidiasis-driven OSCC progression events.

### Oral candidiasis increases p63 and vimentin expression and decreases E-cadherin expression in OSCC histopathological specimens

p63 expression is a reliable indicator in histological grading and is an early marker of poor OSCC prognosis. When comparing the p63 histopathological staining of the OSCC xenograft and OS-OSCC xenograft samples, analysis of OC-OSCC xenograft samples showed that *C. albicans* treatment increased p63 expression and localisation to the nucleus compared to the uninfected tissues (Figure 5A). Next, we analyzed the expression of EMT markers by performing E-cadherin and vimentin staining. Robust expression of E-cadherin was observed in the cell membrane of the OSCC xenograft tumors whereas OC-OSCC xenograft samples demonstrated attenuated E-cadherin expression (Figure 5B). Vimentin was detected in the cytoplasm of only a few cells in OSCC xenograft sections, but many vimentin-positive cells were observed in the OC-OSCC xenograft samples (Figure 5C).

**Figure 5.**
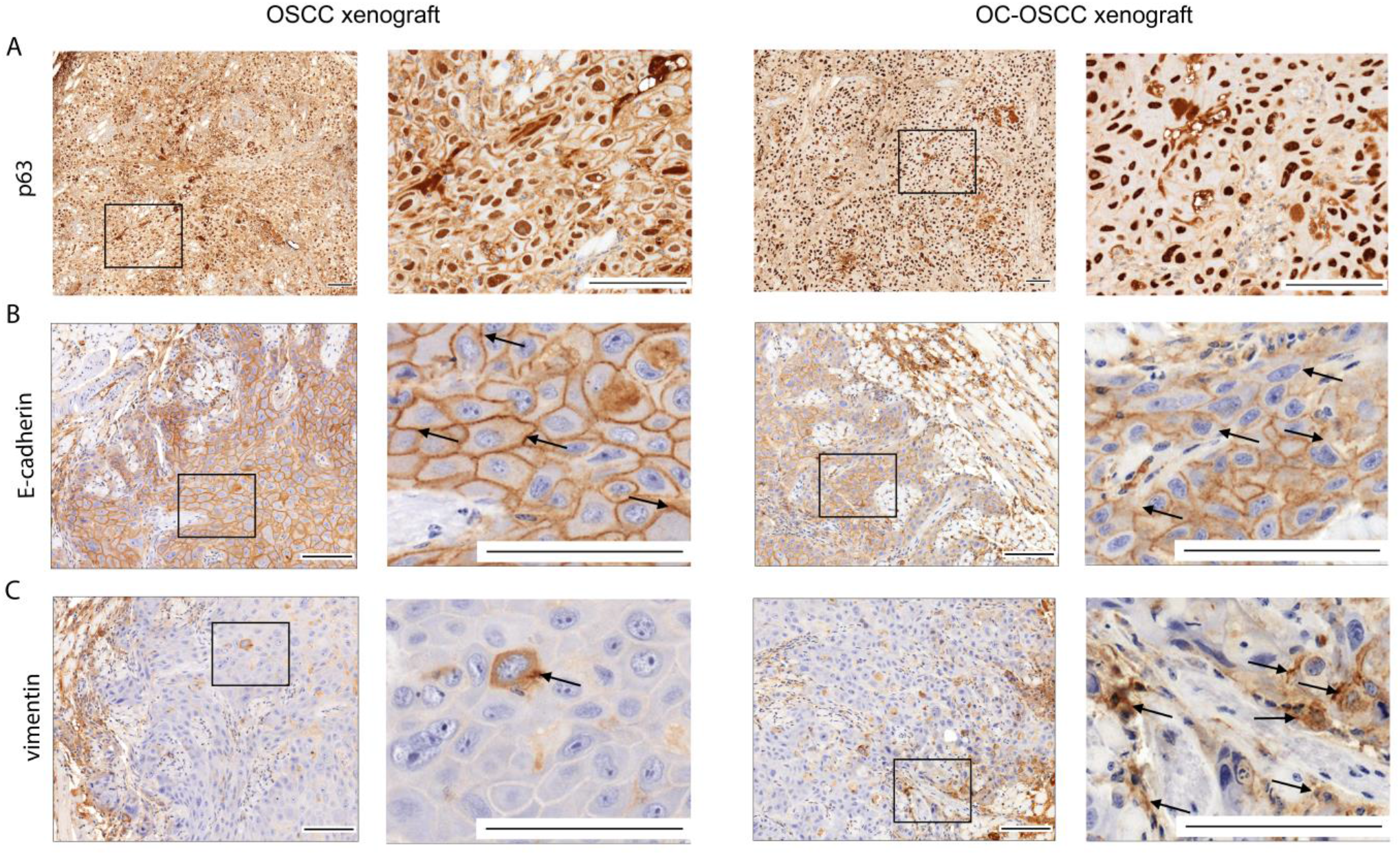
Histopathological staining of OSCC and OC-OSCC xenograft tumor samples: (**A**) p63 staining, (**B**) E-cadherin staining, and (**C**) Vimentin staining. Scale bars: 100 µm. n=4/animal group, derived from four different experiments.

### Oral candidiasis enhances the expression of genes involved in OSCC progression *in vivo*

To analyze the molecular mechanisms behind the histopathological results, transcriptome analysis of OC-OSCC xenograft tumor samples was performed and data were compared to that of OSCC xenograft tumors (Figure 6A). Obtained data analyses showed that oral candidiasis in OSCC xenografts altered the expression of 229 genes (144 upregulated and 85 downregulated) (Supplementary table 2). Among these, 5 genes (MMP10, MMP1, SERPINB4, and CRABP2 upregulated; MMP7 downregulated) are predicted to be involved in OSCC invasion processes (Figure 6B), while 3 genes (MMP10, MMP1, and COL5A2 upregulated) are associated with epithelial-mesenchymal transition regulation (Figure 6C). Notably, MMP10 and MMP1 were present in both subsets (Figure 6D) and were also upregulated in both our *in vitro* and *in vivo* models (Figure 6E).

**Figure 6.**
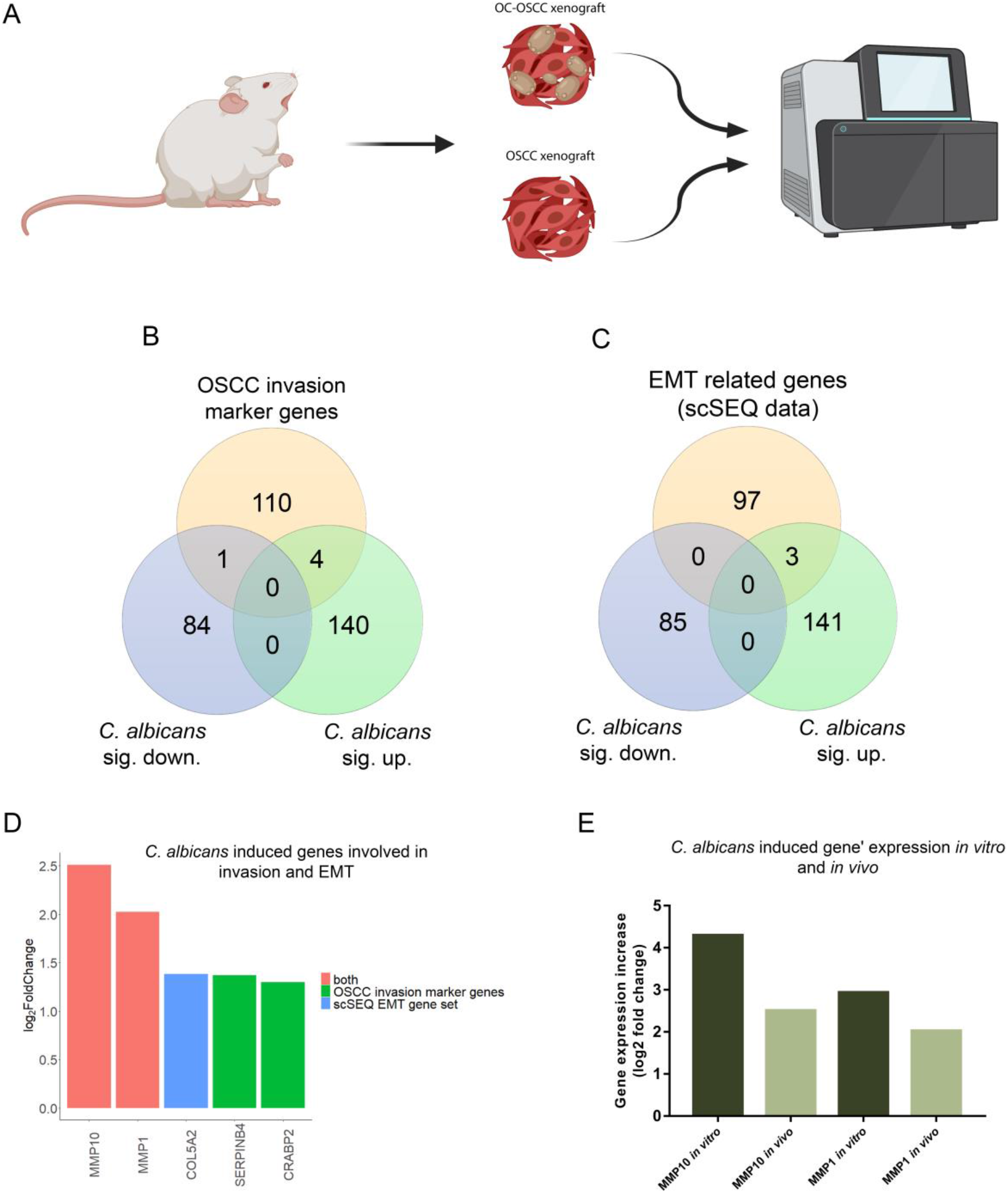
Transcriptomic analysis of *in vivo* tumor samples followed by oral candidiasis. (**A**) Schematic figure of mRNA sequencing of OSCC xenograft and OC-OSCC xenograft tumor samples. The cartoon was produced by BioRender. (**B**) VEN diagram of up or downregulated genes in HSC-2 cells in the presence of live *C. albicans* and OSCC invasion marker genes described in the literature. (**C**) VEN diagram of up or downregulated genes in HSC-2 cell line in the presence of live *C. albicans* and EMT marker genes in HNSCC according to a single cell sequencing study. (**D**) Graph shows log2 fold change of *C. albicans* induced genes in HSC-2 cells involved in OSCC invasion according to the literature and single-cell sequencing data. Red columns show OSCC marker genes according to both the literature and single-cell sequencing (scSeq) results from 18 patients with HNSCC (**E**) Tumor invasion genes showing upregulated expression both *in vitro* and *in vivo*. n= 4/animal group, derived from four different experiments.

## Discussion

OSCC is highly associated with the presence of oral candidiasis. However, the details of the cause versus causative relationship are not understood^25–31^. Our previous study showed that the diversity of the oral fungal microflora of OSCC patients is remarkably different from healthy individuals: the fungal burden and diversity of yeasts was significantly higher in patients with oral tumors compared to the oral cavity of healthy individuals. Furthermore, fungal colonisation in the oral cavity in patients with OSCC is higher on the neoplastic epithelial surface compared to healthy surfaces, which indicates a positive association between oral yeast carriage and epithelial carcinoma ^14^. *Candida* might induce carcinogenesis by the production of carcinogenic compounds, for example, nitrosamines^43^. These carcinogens bind to bases, phosphate residues, and/or hydrogen bonding sites of DNA that could interfere with DNA replication. Induced point mutations might activate oncogenes and initiate the development of oral cancer. Accumulation of acetaldehyde, a by-product of ethanol metabolism that is also considered to be carcinogenic, and the induction of proinflammatory cytokines could also contribute to oral cancer development^44^, both of which are triggered by the presence of *Candida* cells.

*C. albicans* and *C. parapsilosis* both are common commensals of the oral cavity ^45^. Both are also opportunistic human pathogenic fungi, although *C. albicans* is more frequently associated with oral candidiasis compared to *C. parapsilosis*^46^. *C. albicans* is truly polymorphic, due to its ability to form hyphae and/or pseudohypha^47^. Concerning *C. parapsilosis*, this species does not produce true hyphae but can generate pseudohyphae that are characteristically large and curved^48^. Hypha formation is critical for host cell damage and immune activation, which are both driven by the secretion of Candidalysin, a peptide toxin. Candidalysin damages epithelial membranes and activates danger response pathways, which results in immune activation and the secretion of cytokines and chemokines^49^.

In this study, we aimed to examine the potential effects of an increased fungal burden on the progression of oral squamous cell carcinoma. During the experiments, HI-*Candida*, zymosan and live *Candida* cells were applied to two OSCC cell lines, HSC-2 and HO-1-N-1. To examine the potential pro-tumor effect of direct cell-cell interactions, first host cell proliferation and migration were assessed. A previous study showed that Gram-negative bacteria facilitated proliferation and invasion activity of non-small cell lung cancer cells^50^. During our experiments we could not detect any significant changes in host cell proliferation after the applied stimuli; however, the migration activity of the HO-1-N-1 cell line significantly increased after HI-fungal and zymosan treatment. Live cell imaging revealed separately migrating HSC-2 tumor cells from the edges of the disrupted regions of the tumor monolayer after exposure to live *C. albicans* cells. As several of the segregated cells appeared distinctly from the plate-attached hyphae of *C. albicans* cells, we assume that an indirect interaction could also take place between tumor and yeast cells possibly through secreted molecules or extracellular vesicles^51^. This may play a role in tumor progression by the enhancement of invasion. No such phenomenon was observed in the presence of live *C. parapsilosis* cells.

Matrix metalloproteinases (MMPs) are essential to the tumor metastasis processes, as their secretion degrades components of the extracellular matrix elements, facilitating the migration of individual malignantly transformed cells^52^. Notably, as a result of HI fungal challenge and zymosan treatment, we detected a slight increase in secreted MMP activity. Additionally, live *C. albicans* treatment resulted in a marked increase in MMPs activity, supporting the hypothesis that *C. albicans* can increase tumor invasion by the activation of MMPs. Live *C. parapsilosis* treatment did not trigger an increase in MMP concentrations. Our transcriptomic results also support this finding as *C. albicans* induced the expression of several MMP genes in the HSC-2 cell line.

Metastatic tumor cells have unique metabolic profiles^53^. For this reason, we examined the concentrations of glycolysis, TCA cycle intermediated and some amino acids with HPLC-HRMS with and without exposure to *Candida* or zymosan. Metabolic analysis revealed that HI-*Candida* and zymosan treatments result in only a slight and inconsistent alteration in the production of a few metabolites. In contrast, live *C. albicans* cells increased the amount of aspartic acid and succinic acid and decreased the amount of glyceraldehyde-3P in both OSCC cell lines. Transcriptomic and qPCR results also support these findings. Gene expression of GOT1, DLST and SUCLA2, involved in aspartic and succinic acid synthesis, was significantly increased. Interestingly, the amount of glyceraldehyde-3P also decreased in both cell lines after HI-*C. albicans* treatment compared to the untreated control. Succinic and aspartic acid play a role in tumor invasion. Succinate may function as a driver of tumorigenesis through multiple mechanisms. The accumulated succinate itself can inhibit prolyl hydroxylase. This enzyme is responsible for hydroxylation of HIF1α causing its degradation. Therefore, succinate accumulation through inhibition of prolyl hydroxylase causes HIF1α stabilization and its translocation to the nucleus, which might enhance angiogenesis, resistance against apoptosis, and the activation of genes involved in tumor invasion^54^. According to another study, the secreted tumor-derived succinate belongs to a novel class of cancer progression factors, inhibiting tumor-associated macrophage polarization and promoting tumorigenic signaling through PI3K/AKT and HIF-1α^55^. Another study showed that aspartate is a limiting metabolite for cancer cell proliferation under hypoxia and in tumors, such that aspartate might be a limiting metabolite for tumor growth^56^. This suggests that the *C. albicans* induced aspartic acid increase may facilitate OSCC progression. We assume that the reduced GA-3P levels could be the result of increased glyceraldehyde-3-phosphate dehydrogenase (GAPDH) activity. GAPDH catalyzes the redox reaction in the glycolytic pathway by converting GA-3P to 1,3-bisphosphoglycerate with a reduction of NAD^+^ to NADH. GAPDH promotes cancer growth and metastasis through upregulation of SNAIL expression^57^. Increased succinic and aspartic acid, and decreased GA-3P suggest that the presence of live *C. albicans* might enhance OSCC progression by altering complex metabolic processes in tumor cells.

Transcriptome analysis results also supported our *in vitro* findings as the presence of live *C. albicans* cells alters the metastatic features of OSCC cells, while HI-*Candida* and zymosan stimuli had a mild effect. Comparing the two cell lines, HSC-2 showed more prominent responses to live *C. albicans* treatment in terms of the activation of genes and pathways than HO-1-N-1 cells. In both cell lines, we identified several *C. albicans*-induced genes that are involved in OSCC invasion and metastasis regulation (Supplementary table4). Among the overlapping genes, there were 14 *C. albicans*-induced genes (ATF3, F3, FOS, FOXC2, HBEGF, IL6, INHBA, JUN, LIF, PHLDA1, PLAUR, PTHLH, SEMA7A, and VEGFA) whose expression increased in both cell lines.

The obtained transcriptomic data support our *in vitro* findings on total MMP activity and oncometabolite secretion. We detected a significant increase in MMP1, MMP10, MMP3, and MMP9 expression, and a slight increase in the expression of ASNSD1, GOT1, DLST, SUCLA2, all of which are involved in aspartic and succinic acid metabolism. All obtained transcriptomics data were further validated by qPCR.

To examine whether live *C. albicans* enhances invasion and metastatic activity of OSCC *in vivo*, we developed a novel *in vivo* mouse model of OSCC and oral candidiasis. Human HSC-2 OSCC cells were injected into the tongue of the immunosuppressed BALB/c mice. After oral tumor development, xenograft mice were infected with *C. albicans* to induce oral candidiasis development. During the subsequent experiments, yeast-free tumor samples were compared to *C. albicans*-colonized tumors derived from the xenograft groups. In the OC-OSCC tongues, infiltrating immune cells were observed on the mucosa, which was further characterized as having severe inflammation caused by *C. albicans*. Signs of inflammation could also be observed in the tumor tissue. Inflammatory cells promote the development, advancement, and metastasis of cancer by producing tumor-promoting cytokines. Furthermore, inflammation can alter the tumor microenvironment by inducing growth, survival, proangiogenic factors, and reactive oxygen species. It can also modify the extracellular matrix, thereby promoting angiogenesis, invasion, and metastasis^58^. EMT plays a vital role in invasion and metastasis of cancer cells^59^. Thrombosis was also detected in 3 OC-OSCC xenograft samples. The relationship between thrombosis and tumor invasion is well established, but the details of the exact pathological processes are not yet known. Although the details of the process are unknown, thrombosis promotes metastasis^60^. To further evaluate the metastasis enhancing effect of *C. albicans*, we performed p63 staining on histopathological samples. p63 is the protein encoded by the *TP63* gene, a *TP53* gene homolog, known for its key role in cell cycle regulation and in tumor differentiation. Its overexpression is associated with a poor prognosis of head and neck squamous cell carcinoma^61^. p63 has two different promoter domains that produce TAp63, which includes an NH_2_-terminal transactivation domain, and ΔNp63, which lacks an NH_2_-terminal domain. ΔNp63 enhances EMT events during tumor progression by competing with TAp63 and p53 for binding sites^61,62^. In OC-OSCC xenograft samples, p63 expression and localisation to the nucleus was higher compared to the OSCC xenograft samples. We validated this by qPCR in HSC-2 cells in which the ΔNp63 (lacks N-terminal domain) transcript variant can be found (Supplementary 11). In order to further examine EMT progression in OC-OSCC xenograft samples, we performed vimentin and E-cadherin staining. Vimentin is a cytoskeletal protein that is expressed in mesenchymal cells (fibroblast, endothelial cell, lymphocytes), but not in healthy epithelial cells and its upregulation in tumors is linked with lymph node metastasis^63^. Increased vimentin expression is associated with poor prognosis in OSCC^64–66^. In histopathological sections OC-OSCC samples, we detected more vimentin-positive cells than in OSCC samples. E-cadherin, a 120 kDa transmembrane adhesion receptor, apart from being responsible for cell-cell adhesion in the epithelium, plays an important role in determining cell polarity and is also involved in several signal transduction pathways (e.g., induction of apoptosis, growth factor receptor activation)^67,68^. A previous study showed that the reduced expression of E-cadherin is a reliable indicator of increased invasiveness of OSCCs^69^. OSCC tumor samples showed E-cadherin membrane positivity, in contrast to this OC-OSCC samples showed reduced E-cadherin membrane positivity. Increased p63 and vimentin expression and decreased E-cadherin expression in the isolated *Candida* colonized tumors altogether indicate a poor prognosis for oral candidiasis-associated OSCC. Importantly, subsequent transcriptome analysis results of the *in vivo* OC-OSCC samples revealed that the presence of *C. albicans* cells induced a similar effect under these experimental arrangements as in the *in vitro* setting. The expression of MMP1, MMP10, COL5A2, SERPINB4, and CRABP2 was increased under both conditions, confirming the OSCC progression enhancing effect of live *C. albicans* cells.

Oral candidiasis in OSCC patients frequently develops and is thought to exacerbate poor prognosis. Prior to our work, no studies rigorously investigated this phenomenon. In this study we have shown that oral candidiasis indeed aids tumor progression by exacerbating the expression of tumor invasion and metastasis-associated prognostic markers. Given that oral candidiasis frequently develops in oral tumor patients, as the consequence of chemoradiotherapy, it therefore increases the risk for tumor invasion and metastasis. Thus, applying antifungal treatment simultaneously with chemotherapy might be recommended in this patient cohort. Although the efficacy of such a combined treatment option still needs to be addressed, the hereby gained data provide a strong basic understanding of the nature of yeast-tumor interactions which could contribute to the initiation of subsequent translational investigations.

## Materials and methods

### Ethics Statement

The study conformed to EU Directive 2010/63/EU and was approved by the regional Station for Animal Health and Food Control (Csongrád-Csanád, Hungary) under project license XXIX./4061/2020.

### Cell lines and maintenance

Two human oral squamous cell carcinoma (OSCC) cell lines were used. HSC-2 (JCRB0622) cells were cultured in EMEM (Eagle’s Minimum Essential Medium; Lonza), while HO-1-N-1 (JCRB0831) cells were cultured in DMEM/F-12 (Dulbecco’s Modified Eagle Medium:F12; Lonza) medium, both containing 10% heat-inactivated (56°C, 30 min) fetal bovine serum (FBS) (EuroClone) supplemented with 4 mM glutamine, 100 U/mL penicillin and 100 mg/mL streptomycin. Cells were maintained at 37°C in the presence of 5% CO_2_.

### Wound Healing Assay to assess cellular migration

HSC-2 and HO-1-N-1 cells were seeded at 3×10^5^ cells/well in 6-well plates per well. Cells were allowed to grow until 100% confluency. Scratches were made in across the confluent cells using a P100 pipette tip. Zymosan (10 µg/mL working concentration) or heat-killed (HI) (65°C, 2 h) or live *C. albicans* SC5314 (SZMC 1523) or *C. parapsilosis* CLIB 214 (SZMC 1560) were used as fungal treatments. In the case of HI-*Candida*, the multiplicity of infection (MOI) was 1:10 (tumor cell : yeast). Images were taken at time point 0 h (immediately after treatment) and 24 h. Cell migration speed was analyzed using imageJ software.

The rapid hyphae formation of live *C. albicans* cells did not allow a comprehensive analysis of the extent of cancer cell movement after 24 h. Thus, a 24 h time-lapse (CytoSMART) video was analysed to examine the extent of tumor cell migration. For this, experiment, 1×10^5^ tumor cells were seeded in to 24-well plates using 500 µm culture inserts and grown until full confluency. On the following day, the insert was removed and live *Candida* (*C. albicans* MOI 1:400; *C. parapsilosis* MOI 1:4) were added and imaged by time-lapse video with CytoSMART Lux2.

### BrdU incorporation assay

Cell proliferation activity was measured by BrdU incorporation assay using a Cell Proliferation ELISA kit (Sigma-Aldrich). The wells of 96-well plates were seeded with 5000 HSC-2 or HO-1-N-1 cells. The following day, cells were treated with zymosan (10 µg/mL), HI-*Candida* (MOI 1:10), live *C. albicans* (MOI 1:400) or *C. parapsilosis* (MOI 1:4) for 24h, and then BrdU assay was performed according to the manufacturer’s instructions. The experiment was performed in media supplemented with 1% FBS, 4 mM glutamine, 100 U/mL penicillin and 100 mg/mL streptomycin.

### Sample preparation for metabolic analysis

For metabolomic analyses, 1.5×10^5^ HSC-2 and HO-1-N-1 tumor cells were seeded per well in 6-well plates. On the following day, cells were treated with zymosan (10 µg/mL) or HI-*Candida* (MOI 1:10) for 24h. After removal of medium, tumor cells were washed with PBS and extracted by addition of 500 μl ice cold mixture of HPLC grade methanol:water (4:6 v/v), and the remaining debris was removed by scraping (20 mm blade width). Cell lysates was transferred to microcentrifuge tubes and sonicated for 5 min at 23kHz in an ice water bath. Sonicated samples were mixed for 15 sec and centrifuged at 13.800xg for 10 min at 4°C. The supernatants were transferred to HPLC vials and stored at −80°C.

Due to the rapid hyphal formation of the live *C. albicans*, for the live yeast conditions, HSC-2 and HO-1-N-1 cells were seeded in 6-well plates (1.5×10^5^/well) in 5 technical replicates. After 4 h, the completely attached cancer cells were treated with live *C. albicans* (MOI 1:400) or *C. parapsilosis* (MOI 1:4) for 24 h. After removal of culture media, the cells were washed immediatelly 1x in 37°C Ringer’s solution, than 2x in 37°C HPLC grade distilled water. Metabolite extraction was performed by incubation of the samples in 500 μl HPLC grade distilled water for 15 min causing osmotic shock for the tumor cells, while the fungal cell wall remained intact according to the literature^70^ and based on our measurement of the supernatants collected from the *Candida* cells incubated in distilled water for 15 min, where no any metabolites were detected. Samples were then centrifuged at 13.800xg for 10min at 4°C and the supernatants were transferred to HPLC vials and stored at −80°C.

### High performance liquid chromatography coupled with high resolution mass spectrometry (HPLC-HRMS) analysis of metabolites

The amount of the intermediaries of glycolysis, TCA cycle and amino acids were determined by HPLC-HRMS. The measurements were carried out on a Dionex UltiMate 3000 (Thermo Scientific) HPLC system coupled to a Q Exactive Plus (Thermo Scientific) HRMS, where the eluent A was water and eluent B was methanol supplemented both of them with 0.1% acetic acid. The applied gradient program was the following on a Synergi Polar-RP (Phenomenex) 250×3 mm, 4 µm column for the eluent B: 0 min −20%, 2 min – 20%, 4 min – 30%, 6 min – 95%, 9 min −95%, 9.5 min −20% and 15 min 20%. The flow rate and the injection volume were 0.2 mL/min and 5 µL, respectively, while the column and the autosampler were thermostated at 30°C and 4°C, respectively. The HRMS was operated with heated electrospray ionization (HESI) source in parallel reaction monitoring (PRM) acquisition mode with polarity switching. During the measurements, the spray voltages were 4 kV and 3 kV in positive-and negative ionization modes, respectively. The sheath gas was 30, the aux gas was 15 arbitrary unit, the aux gas heater temperature was 250°C and the ion transfer capillary temperature was 250°C in both ionization modes. The isolation window was 0.4 *m/z* and the resolution was 35000 (at *m/z*=200). The precursor mass, fragment ion mass, polarity, retention time and fragmentation energy and the lower limit of determination (LLOQ) of the examined metabolites are detailed in (Supplementary table 1). For quantitative determinations, seven level calibration curves were used in the case of each metabolite, in the range of 5-5000 ng/mL. The calibration standard solutions were created in MeOH:H2O 4:6 solution with equal amount internal standard (250 ng/mL) compared with the tumor cell extracts. For the live *Candida* treatment conditions, calibration standard solutions were created in HPLC grade distilled water. Finally, the concentration values were compared with the control samples.

### Matrix metalloproteinases (MMP) enzymatic activity

MMP activity was measured using the MMP activity Assay Kit (Abcam, ab112146), following the manufacturer’s instructions. For the experiments 3×10^5^ cells were seeded into T25 flasks. On the following day, cells were treated with zymosan (10 µg/mL), HI-*Candida* (MOI 1:10), live*C. albicans* (MOI 1:400) or *C. parapsilosis* (MOI 1:4) for 24 h in 4 mL serum-free medium. Next, the media was collected and centrifuged for 5 min, 3000xg and 4 mL of the supernatant was concentrated to appoximately 200 µL by centrifugation for 25 min at 7500xg using a centrifugal filter (Amicon Ultra-4, UFC800324). The concentrated samples were adjusted to same volume. Activities of MMPs was measured with fluorescence plate reader at Ex/Em = 490/525 nm.

### RNA extraction for sequencing *in vitro* samples

For RNA extraction, 1×10^5^ HSC-2 and HO-1-N-1 cells were seeded into 24-well plates with three technical replicates. After 12 h, the tumor cells were treated with zymosan (10 µg/mL), HI-*C. albicans* (MOI 1:10), HI-*C. parapsilosis* (MOI 1:10), live *C. albicans* (MOI 1:25) or live *C. parapsilosis* (MOI 1:4) for 12 h. After fungal treatment, mRNA was purified using the RNeasy Plus Mini Kit (Qiagen) according to the manufacturer’s protocol. RNA quality and quantity was analyzed by Bioanalyzer Instrument (Agilent). Library preparation and sequencing on a NovaSeq S4 platform was performed by Novogene.

### RNA Sequencing

The preparation of mRNA sequencing library was done by external specialists at Novogene. Briefly, sequencing libraries were generated using NEBNext® Ultra TM RNA Library Prep Kit for Illumina® (NEB, USA) following the manufacturer’s recommendations. For the purification of mRNA samples, poly-T oligo-attached magnetic beads were used, then size selected cDNAs were synthesized with Ampure XP system (Beckman Coulter, Beverly, USA).

### Transcriptome analysis

RNA sequence files were also processed by Novogene; however, the analyses of *in vivo* samples were repeated using our in-house protocol. According to the Novogene analysis pipeline, raw sequence reads processed for quality via fastp, reads with adaptor contamination, high precentage (N > 10%) of misread or uncertan bases and low quality (phred < 20) were filtered out. Clean read files were aligned to the reference genome index (GRCh38) using ‘HISAT2’, with the parameters --dta –phred33. Read counts of known and novel genes were quantified as reads per kilobase of exon model per Million mapped reads (RPKM) via ‘Featurecounts’. This step in the case of the *in vivo* samples was interchanged with the ‘GenomicAlignments’ package resulting in counts. Differential gene expression in logarithmic fold change (LFC) was then concluded using the ‘DeSeq2’ tool. Objects with read counts lower, than 1 part per million (ppm) were filtered out. In the experimentally derived gene list, differentially expressed genes (DEGs) were assigned above the absolute value of LFC > 1 and the adjusted p-value < 0.05. False discovery rate was minimized using the Benjamini and Hochberg’s approach.

### Causal analyses

We employed causal analyses methods included in Ingenuity Pathway Analysis (IPA), including (i) Upstream regulatory analysis (URA) to identify probable upstream regulators and (ii) Causal network analysis (CNA) to observe connections between these above mentioned regulatory molecules. A priori an expression core analysis was run on the experimentally derived gene set to obtain a suitable input for these analysis, and predictions with p-value of overlap < 0.05 and z-score different than 0 were considered significant hits. We further specified that only experimentally proven or strongly predicted intermolecular relationships should be considered.

### cDNA synthesis and reverse transcription-PCR for validation sequencing data

A total of 1 µg RNA was used for cDNA synthesis using a RevertAid First Strand cDNA synthesis kit (Thermo Scientific) according to the manufacturer’s instructions. Real-time PCR was carried out in a final volume of 20 μl using Maxima SYBR green/fluorescein qPCR (2x) Master Mix (Thermo Scientific). The reaction was performed in a C1000 thermal cycler (Bio-Rad) using the following reaction conditions: 95°C for 3 min, 95°C for 10 s, 60°C for 30 s, and 65°C for 5 s for 50 cycles. Fold change in mRNA expression was calculated by the threshold cycle (ΔΔCT) method (real-time PCR applications guide; Bio-Rad) using *B2m* housekeeping gene as an internal control.

### Establishment of a novel *in vivo* mouse xenograft model of OSCC and oral candidiasis

6-8 weeks old female BALB/c mice were immuncompromised with cortisone acetate for subsequent injection with human HSC-2 cells. Cortisone acetate was suspended in sterile Ringer’s solution containing 0.05% Tween 80 (v/v) and administered daily *per os* at a concentration of 225 mg kg^-1 42^ in a total volume of 0.2 mL using a sterile gavage (38×22G curved). Because of the immunosuppression, autoclaved rodent feed and bedding were used. Drinking water was supplemented with 1% 100 U/mL penicillin and 100 mg/mL streptomycin. Tumor cell injection was performed on the second day. HSC-2 cells were washed twice with 1xPBS and trypsinized. Trypsin was then neutralized with complete growth medium, and the cell suspension was centrifuged at 400xg for 5 min. After removing the supernatant, the cell pellet was suspended in serum free medium. Anesthesia was performed by administration of 40 mg/kg pentobarbital i.p. and 1×10^6^ HSC-2 cells were injected to the apex of the tongue in 50 µL volume containing 10% (v/v) matrigel (Corning). Mice were continuously monitored until they recovered from anesthesia.

Oral candidiasis was induced on the 5^th^ day with minor modifications to the protocol described previously by Solis et. al^42^. Briefly, *C. albicans* (SC5314) cells were inoculated in 5 mL YPD and cultured for 24 h with continuous shaking at 30°C overnight. Inoculation and culturing was performed 3 times. *Candida* cells were then washed 3 times in 1xPBS, and suspended in HBSS to a concentration of 1×10^9^ cells mL^-1^. The *C. albicans* suspension was placed in 30°C water bath for 5 min. Calcium alginate swabs were placed in the suspension approximately 5 min prior to use. Anesthesia was performed by administration of 40 mg/kg pentobarbital i.p. Saturated calcium alginate swabs were placed under the tongues of each mouse for 75 minutes. The following day, cortisone acetate was administered intraperitoneally to avoid damage in the mouth. No cortisone was given at 7^th^ and 8^th^ day beacause cortisone acetate absorbes slower through i.p. injection resulting prolonged immunosuppressive effect. The weight of the animals was monitored every day. For validation of oral candidiasis, the tongue was excised and homogenized for approximately 8–10 sec in PBS with a tissue homogenizer. Homogenates were used for determination of fungal burdens by colony counting after plating serial dilutions on YPD agar plates per tissue. The CFUs were counted after 48 h of incubation at 30°C and expressed as CFU/g tissue.

### Sample preparation for sequencing from *in vivo* samples

At the last day (8^th^) of the *in vivo* experiment mice were sacrificed and tongues were removed for subsequent analyses. Mouse tongue tissues were removed from the tumor with the help of scalpel. Next, 30 mg tumor tissue was homogenized (Bioneer) in RLT buffer and RNA was extracted with RNeasy Plus Mini Kit (Qiagen) according to the manufacturer’s instructions. RNA quality and quantity were analyzed by Bioanalyzer Instrument (Agilent).

### Histopathological Analysis

Mice were sacrificed and tongues were removed from the base by dissecting scissors and forceps. Whole tongues were fixed in 4% formalin and kept at room temperature until specimen analysis. Fixed tongues containing tumor were sectioned and stained with periodic acid-Schiff (PAS) and hematoxylin-eosin (H&E) using conventional staining methods. For E-cadherin staining, standard immunohistochemical procedures were applied using rabbit monoclonal primary antibody (dilution: 1:200, EP700Y, Cellmarque). For vimentin staining, rabbit monoclonal primary antibody (dilution: 1:300, SP20, Cellmarque) was used. For p63 staining, mouse monoclonal primary antibody (dilution: 1:100, DBR16.1, Hisztopatológia Kft., Pécs, Hungary) was used. Antigen retrieval was performed by epitope retrieval (ER) 2 solution (Leica, pH 9). The slides were then incubated with anti-E-cadherin, anti-vimentin or anti-p63 antibodies for 20 min. Staining was performed with a Leica Bond Max Automatic Staining System using Bond Polymer Refine Detection (Leica). The slides were mounted with coverslips and assessed by microscopic examination (BX51 OLYMPUS or Zeiss Imager Z1)

### Statistical analysis

Statistical analysis was performed using the GraphPad Prism 7 software. All experiments were performed at least three times. Each replicate was normalized to its own control value, when it was necessary, then normalized data was statistically analyzed. Paired or unpaired t-tests were used to determine statistical significance (see figure legends for details) and differences between groups were considered significant at p values of <0.05 (* p ≤ 0.05; ** p ≤ 0.01; *** p ≤ 0.001; **** p ≤ 0.0001).

## ACKNOWLEDGMENTS

This work was supported by 20391-3/2018/FEKUSTRAT; NKFIH K 123952; LP2018-15/2018; ÚNKP-20-5-SZTE-655 (M.K.) New National Excellence Program of the Ministry for Innovation and Technology from the source of the National Research, Development, and Innovation Fund; János Bolyai Research Scholarship of the Hungarian Academy of Sciences (BO/00878/19/8 for M.K.); National Research, Development, and Innovation Office-NKFIH through projects GINOP-2.3.2-15-2016-00038 and GINOP-2.3.2-15-2016-00035 is gratefully acknowledged.

## AUTHOR CONTRIBUTIONS

GA and VM contributed to the concept and design of this project. VM carried out the majority of experiments with the help of IN, RA, RD, VÉ, SZB, HP, TL, VC, SZA, KM, PLG. HM, ASZ analyzed the acquired bioinformatics data. VM prepared the manuscript and the figures, that was revised by RT, JND with AG. All authors reviewed the manuscript, contributed to the discussion and approved the final version.

## DECLARATION OF INTERESTS

The authors declare no competing interests.

Processed and raw expression data is available through the Gene Expression Omnibus (https://www.ncbi.nlm.nih.gov/geo/) with accession number GSE169278.

