## Supplementary figures for "*Candida albicans* enhances the progression of oral squamous cell cancrinoma *in vitro* and *in vivo*"

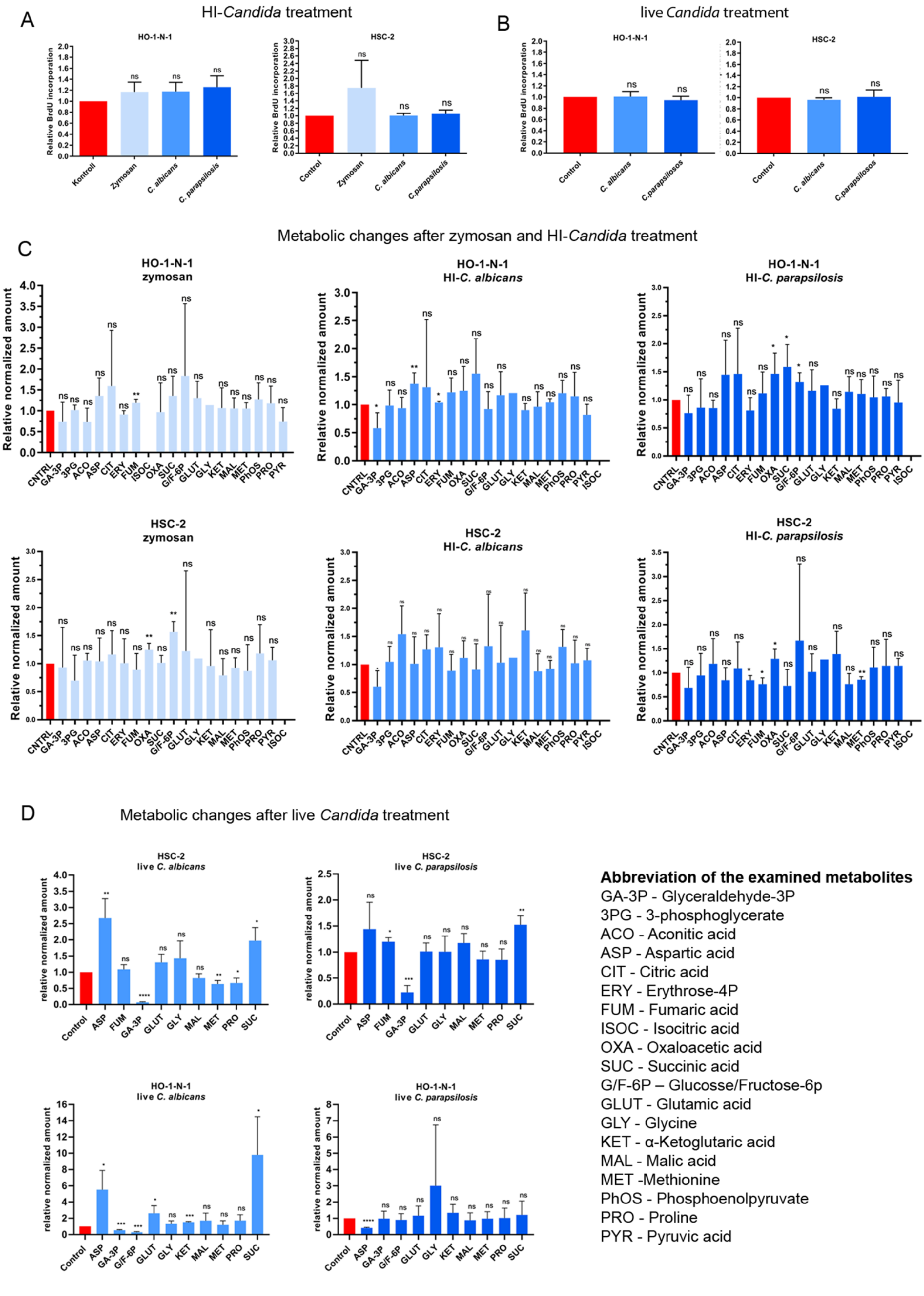
A
HI-Candida treatment
HO-1-N-1
HSC-2
Relative BrdU incorporation
Kontroll
Control
Zymosan
C. albicans
C. parapsilosis
B
live Candida treatment
C.parapsilosos
C
Metabolic changes after zymosan and HI-Candida treatment
HO-1-N-1 zymosan
HO-1-N-1 HI-C. albicans
HO-1-N-1 HI-C. parapsilosis
HSC-2 zymosan
HSC-2 HI-C. albicans
HSC-2 HI-C. parapsilosis
Relative normalized amount
D
Metabolic changes after live Candida treatment
HSC-2 live C. albicans
HSC-2 live C. parapsilosis
HO-1-N-1 live C. albicans
HO-1-N-1 live C. parapsilosis
relative normalized amount
Abbreviation of the examined metabolites
GA-3P - Glyceraldehyde-3P
3PG - 3-phosphoglycerate
ACO - Aconitic acid
ASP - Aspartic acid
CIT - Citric acid
ERY - Erythrose-4P
FUM - Fumaric acid
ISOC - Isocitric acid
OXA - Oxaloacetic acid
SUC - Succinic acid
G/F-6P – Glucosse/Fructose-6p
GLUT - Glutamic acid
GLY - Glycine
KET - α-Ketoglutaric acid
MAL - Malic acid
MET -Methionine
PhOS - Phosphoenolpyruvate
PRO - Proline
PYR - Pyruvic acid

**Supp Fig1**

(A) Normalized proliferation activity of OSCC cells in the presence of heat- killed *C. albicans*, C. *parapsilosis* and zymosan measured by BrdU incorporation assay.

(B) Normalized proliferation activity of OSCC cells in the presence of live *C. albicans* and *C. parapsilosis* measured by BrdU incorporation assay.

(C) Normalized amount of metabolites of OSCC cells in the presence of heat- killed *C. albicans*, *C. parapsilosis* and zymosan measured by HPLC-HRMS.

(D) Normalized amount of metabolites of OSCC cells in the presence of live *C. albicans* and *C. parapsilosis* measured by HPLC-HRMS.

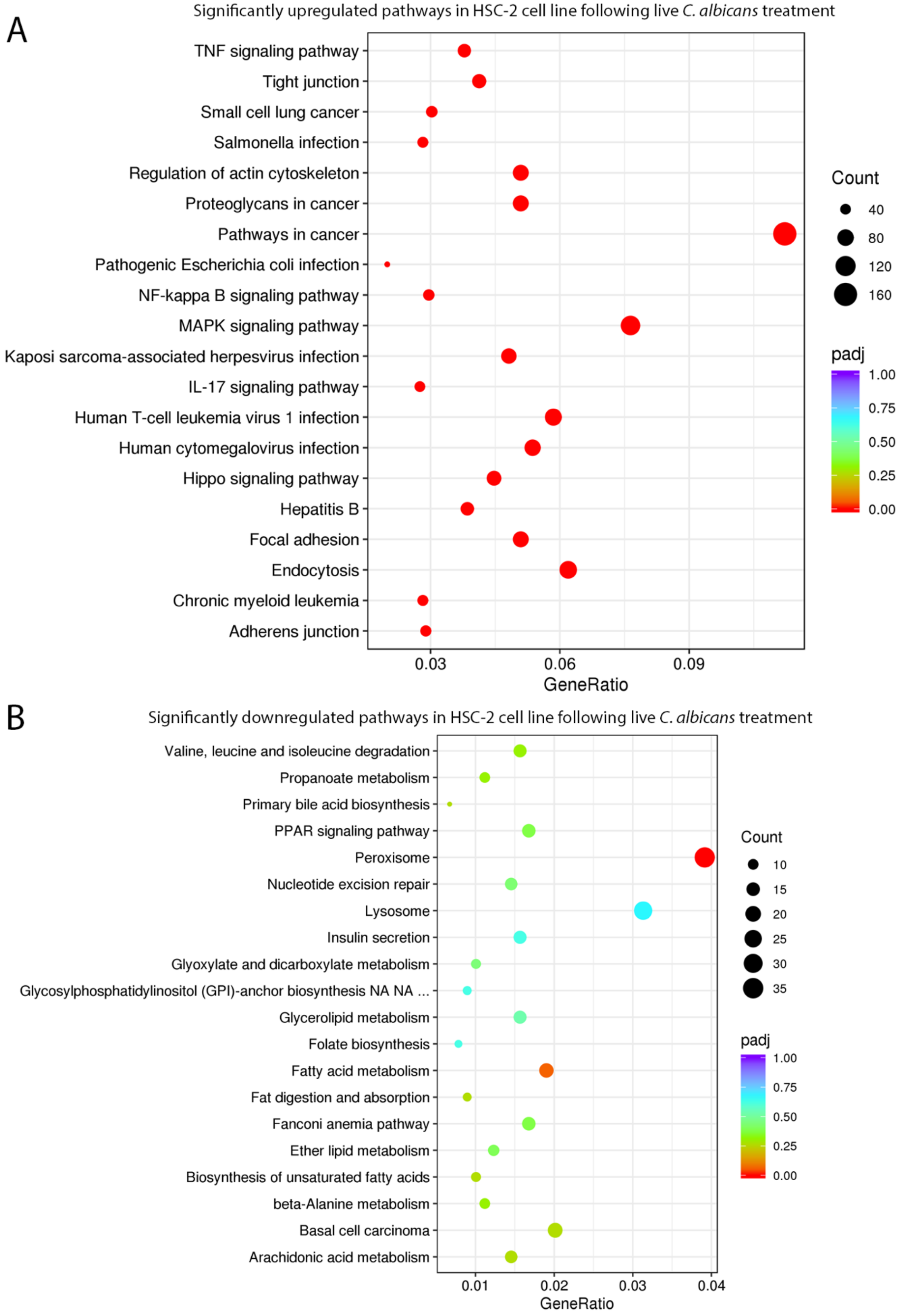


**Supp Fig2**

(A) Significantly upregulated pathways in HSC-2 cell line induced by *C. albicans*

(B) Significantly downregulated pathways in HSC-2 cell line following *C. albicans* treatment

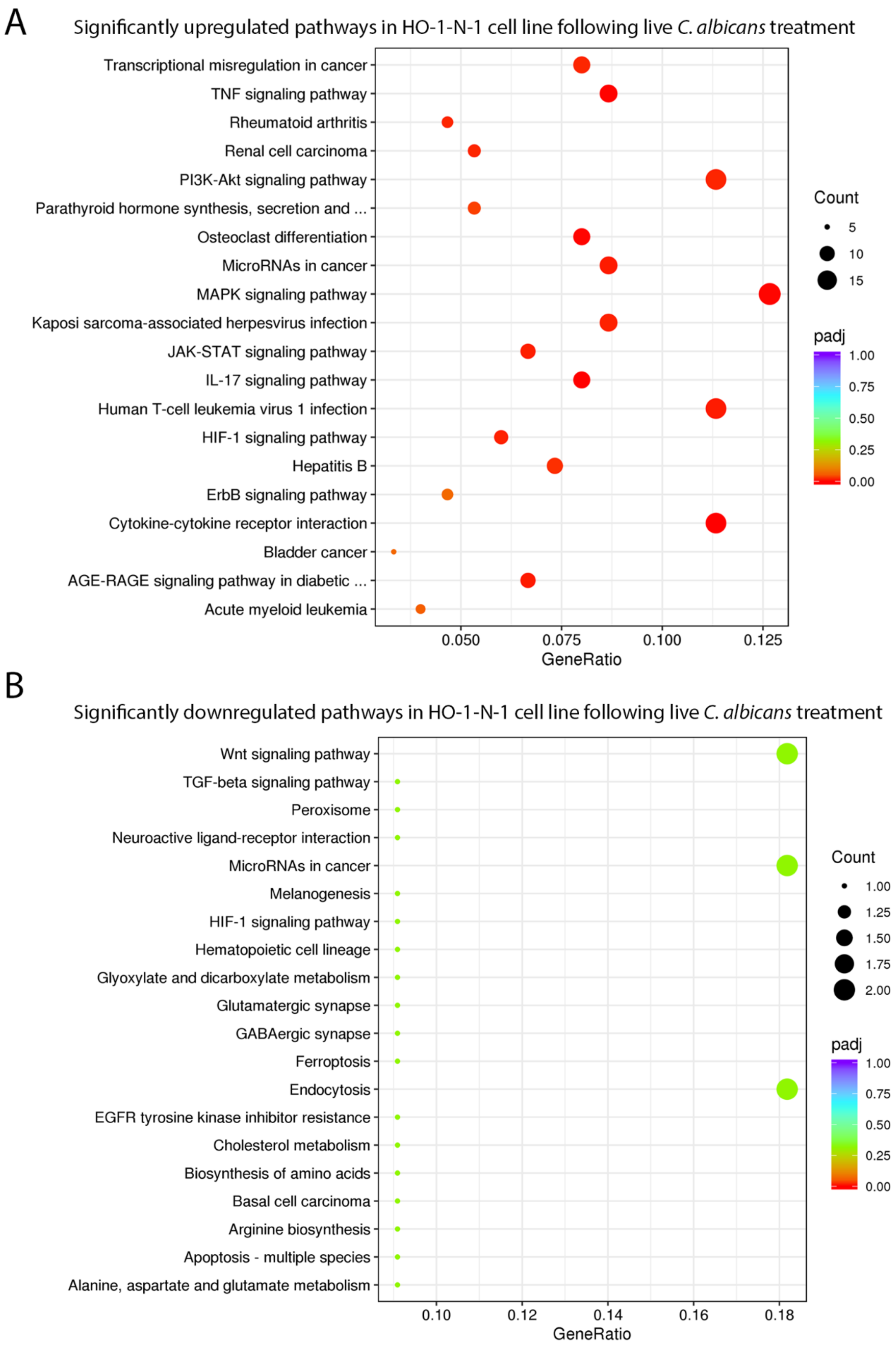


**Supp Fig3**

(A) Significantly upregulated pathways in HO-1-N-1 cell line induced by *C. albicans*

(B) Significantly downregulated pathways in HO-1-N-1 cell line following *C. albicans* treatment

#### Pathways activated in HSC-2 cell line after live *C. albicans* treatment

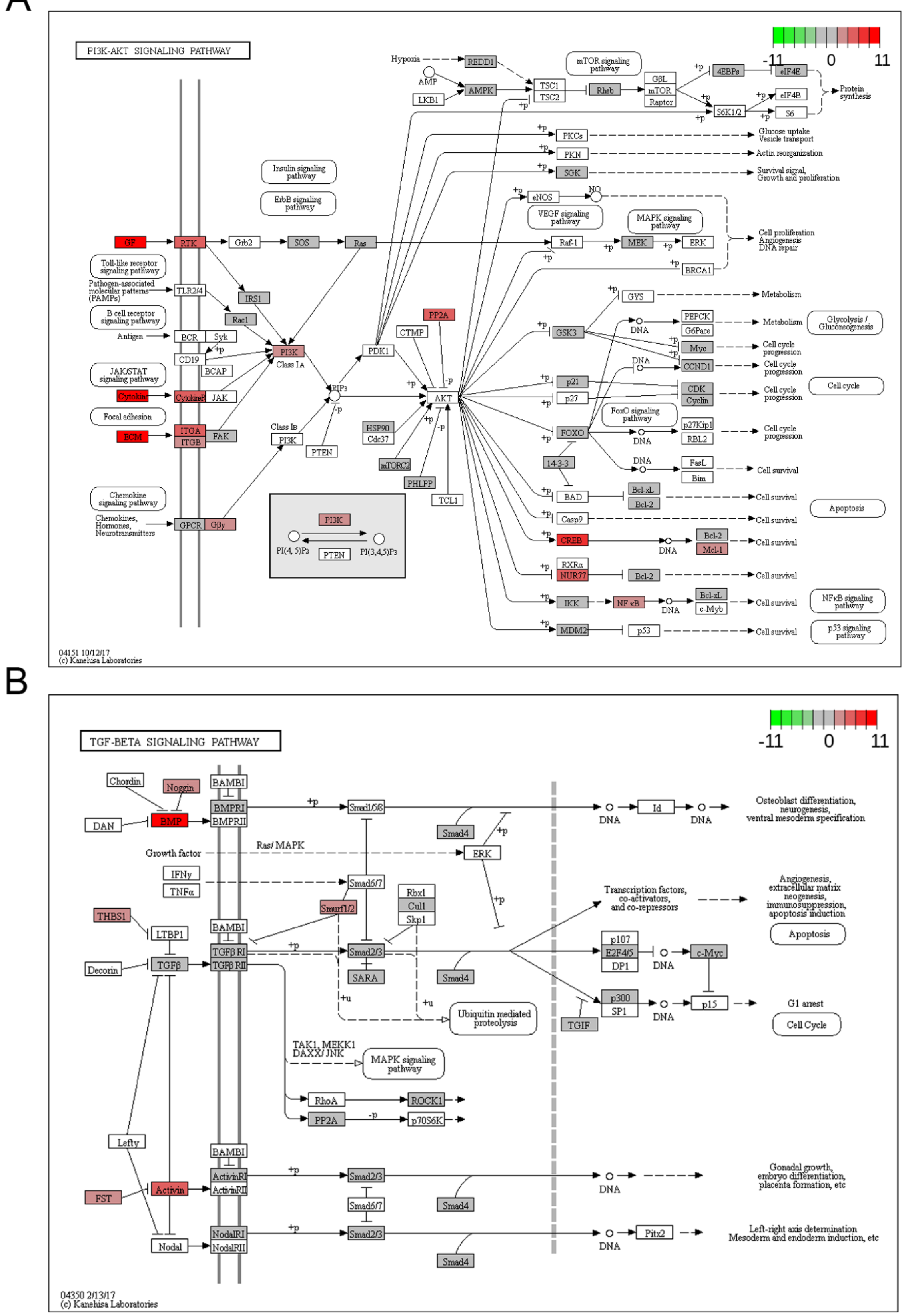


**Supp Fig4**

(A) PI3K-Akt signaling pathway activated in HSC-2 cell line following live *C. albicans* treatment

(B) TGF- β/SMAD signaling pathway activated in HSC-2 cell line following live *C. albicans* treatment

#### Pathways activated in HSC-2 cell line after live *C. albicans* treatment

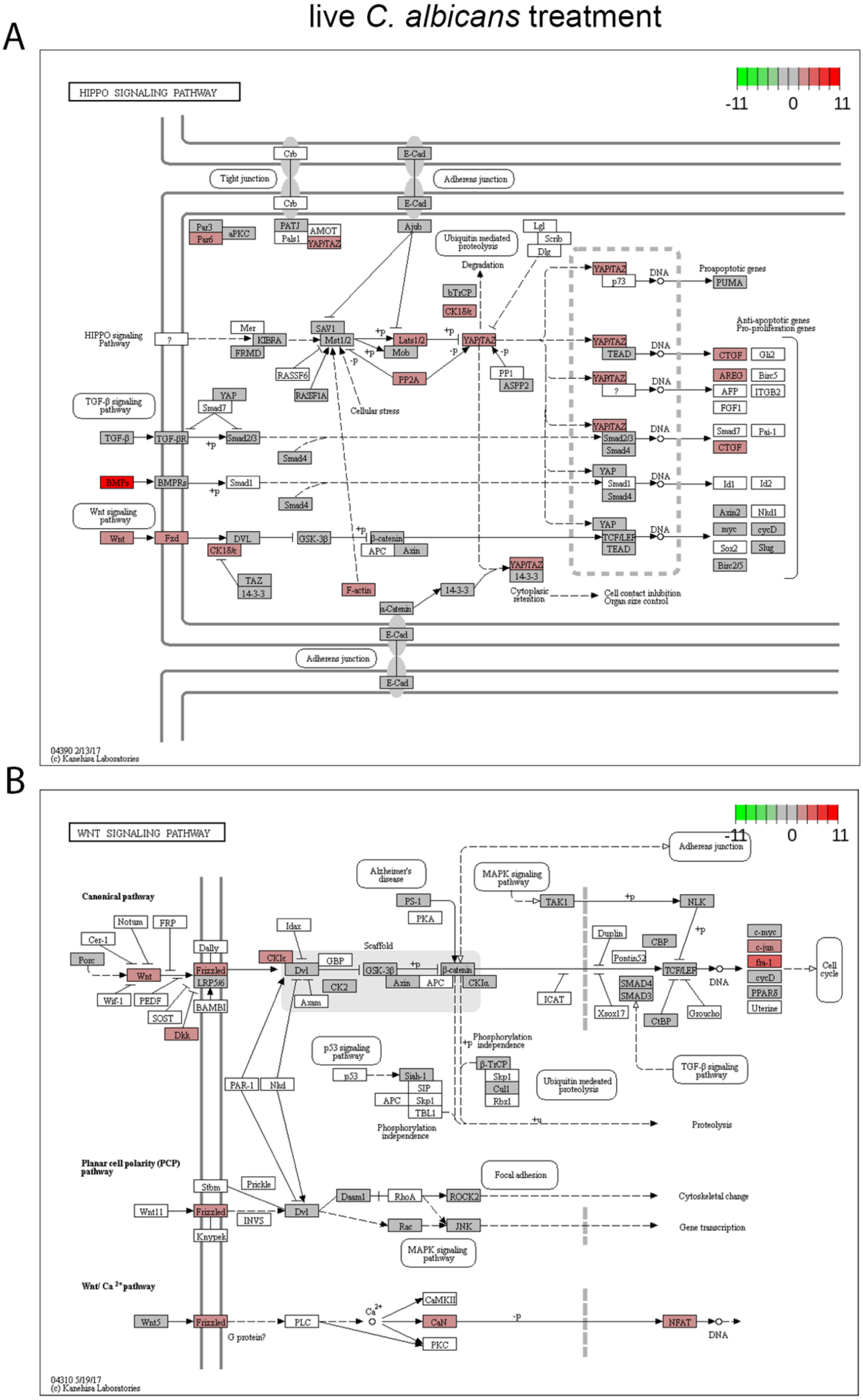


**Supp Fig5**

(A) Hippo signaling pathway activated in HSC-2 cell line following live *C. albicans* treatment

(B) Wnt signaling pathway activated in HSC-2 cell line following live *C. albicans* treatment

Pathways activated in HSC-2 cell line after live *C. albicans* treatment

A

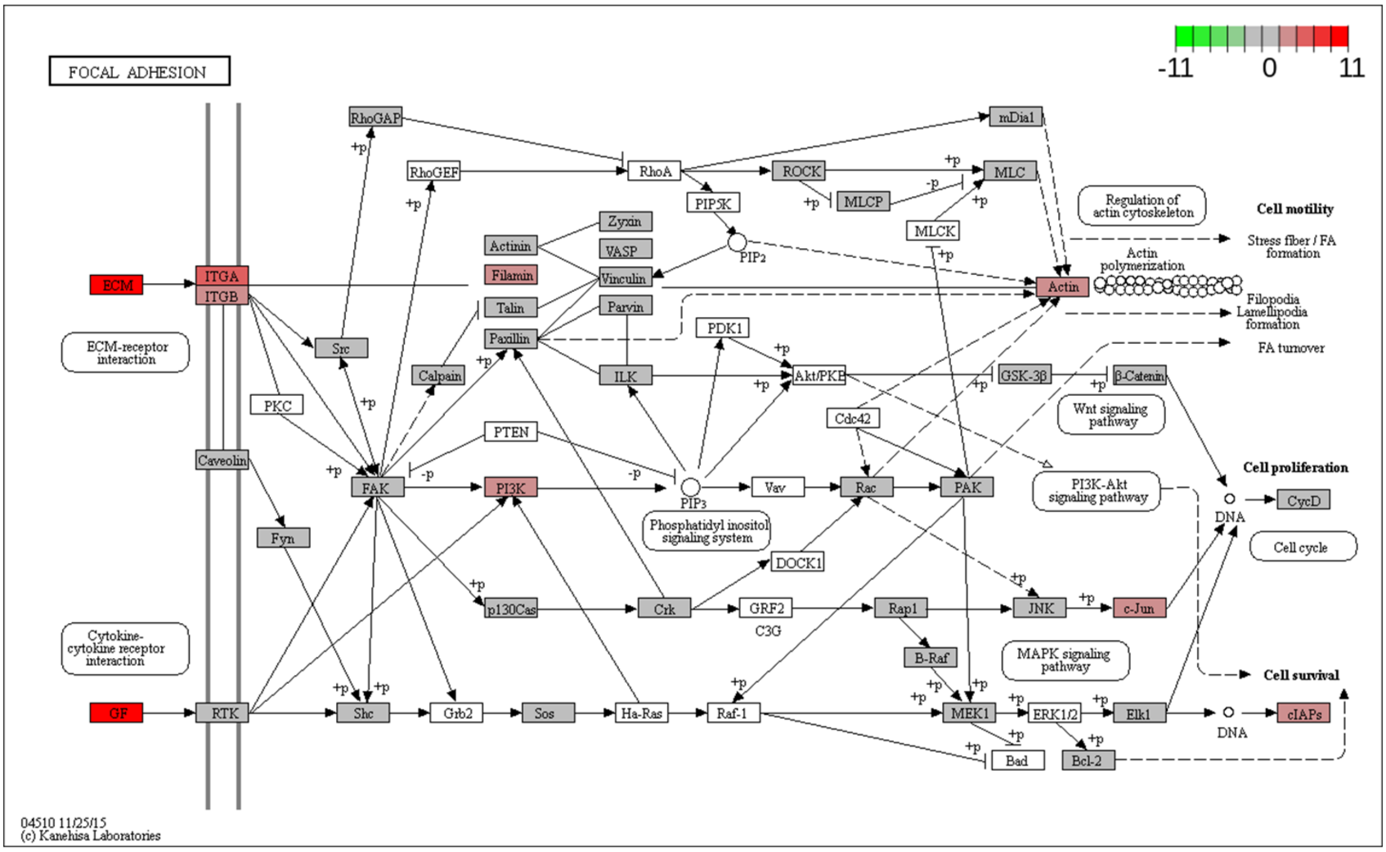


**Supp Fig6**

(A) Focal adhesion pathway activated in HSC-2 cell line following live *C. albicans* treatment

### A: validation of PI3K-Akt pathway components by qPCR

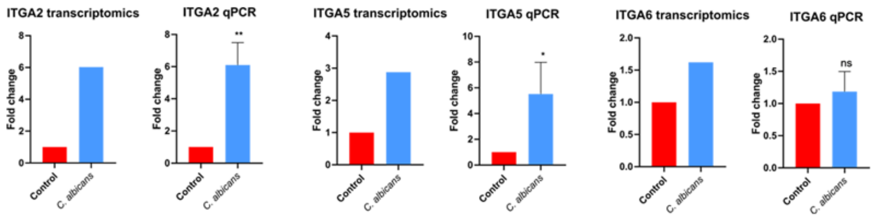


### B: validation of TGFB/SMAD pathway components by qPCR

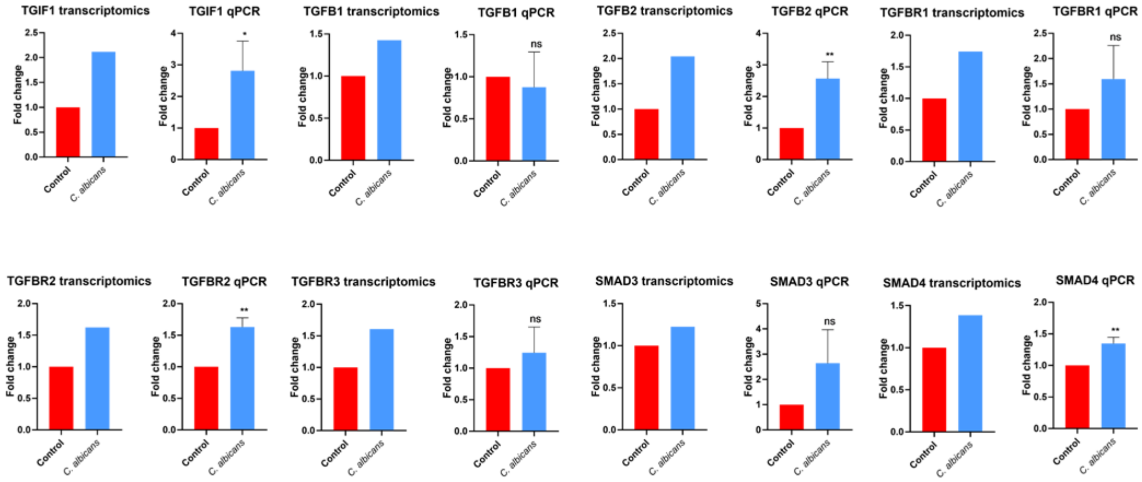


### C: validation of HIPPO pathway components by qPCR

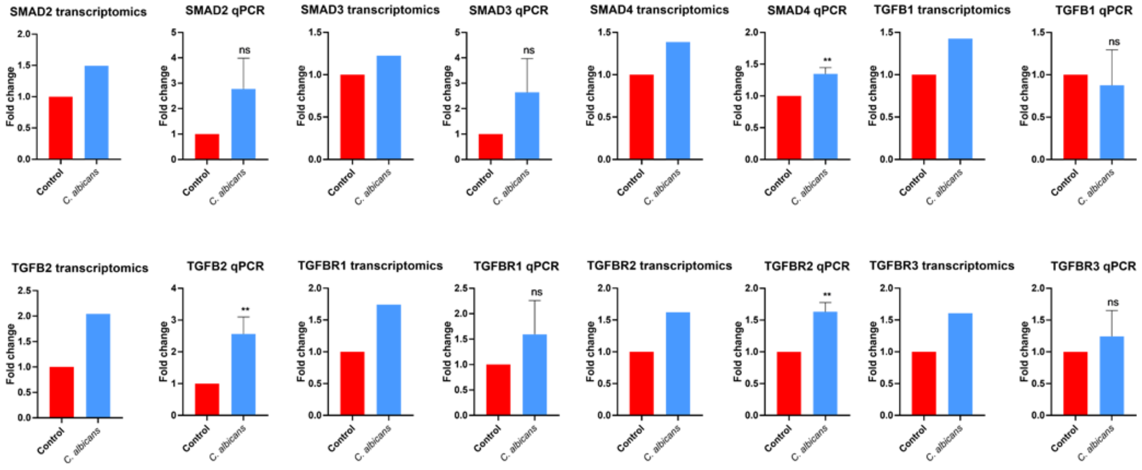


### D: validation of Focal adhesion pathway components by qPCR

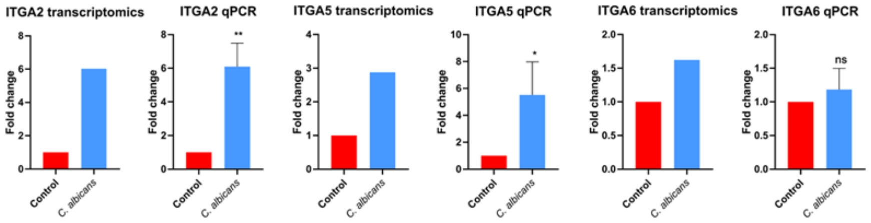


### E: validation of Wnt pathway components by qPCR

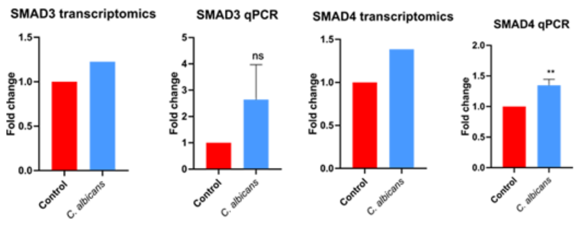

**Supp Fig7**

Validation of *C. albicans* activated signaling pathways. Validation was performed by qPCR analysis of pathway components.

(A) PI3K-Akt signaling pathway (B) TGF- β/SMAD (C) Hippo signaling pathway (D) Wnt signaling pathway (E) Focal adhesion pathway

A HSC-2 cell line *C. albicans* treatment in vitro; scSec data vs. *C. albicans* induced gene set

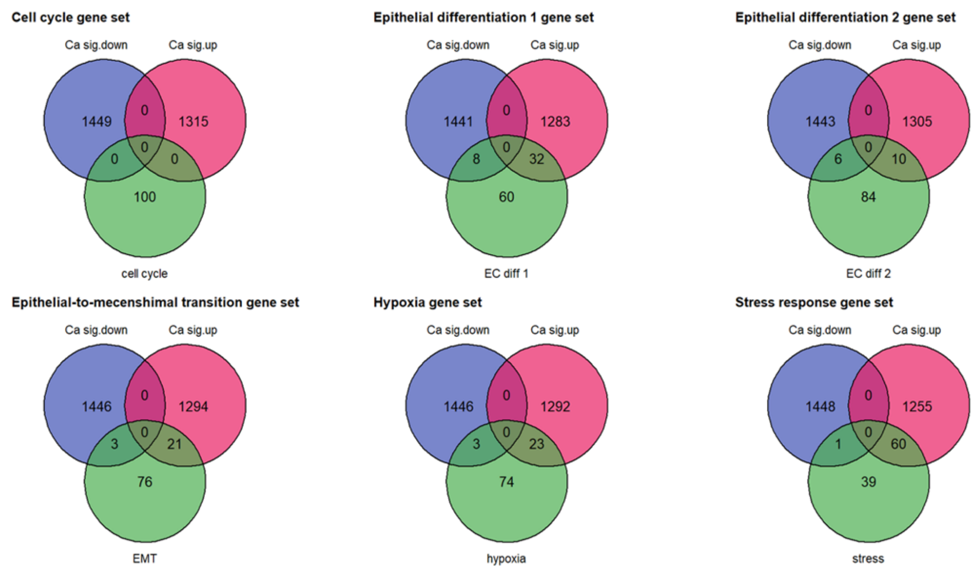


B HO-1-N-1 cell line *C. albicans* treatment in vitro; scSec data vs. *C. albicans* induced gene set

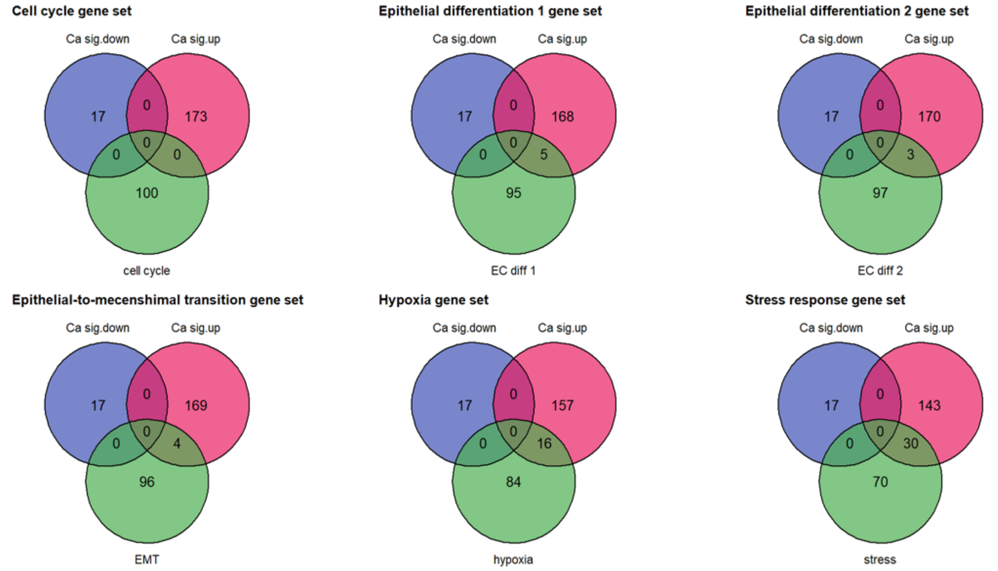


C

Live *Candida albicans* induced genes in both (HSC-2 and HO-1-N-1) cell lines, which are involved in OSCC progression

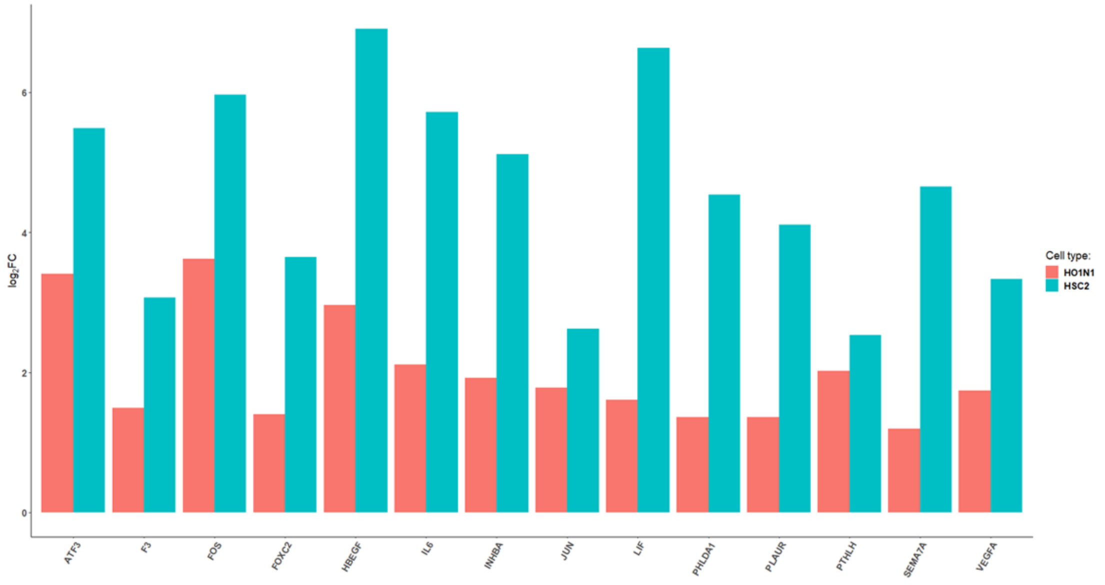

**Supp Fig8**

(A) Comparison of *Candida induced* genes in HSC-2 cell line to genes involved in different tumor progression processes. Differentially expressed gene list derived from an OSCC single cell sequencing study.

(B) Comparison of *Candida induced* genes in HO-1-N-1 cell line to genes involved in different tumor progression processes. Differentially expressed gene list derived from an OSCC single cell sequencing study.

(C) Live *C. albicans* induced genes in both (HSC-2 and HO-1-N-1) cell lines, which are involved in OSCC progression. OSCC progression marker gene list derived from literature and OSCC single cell sequencing study.

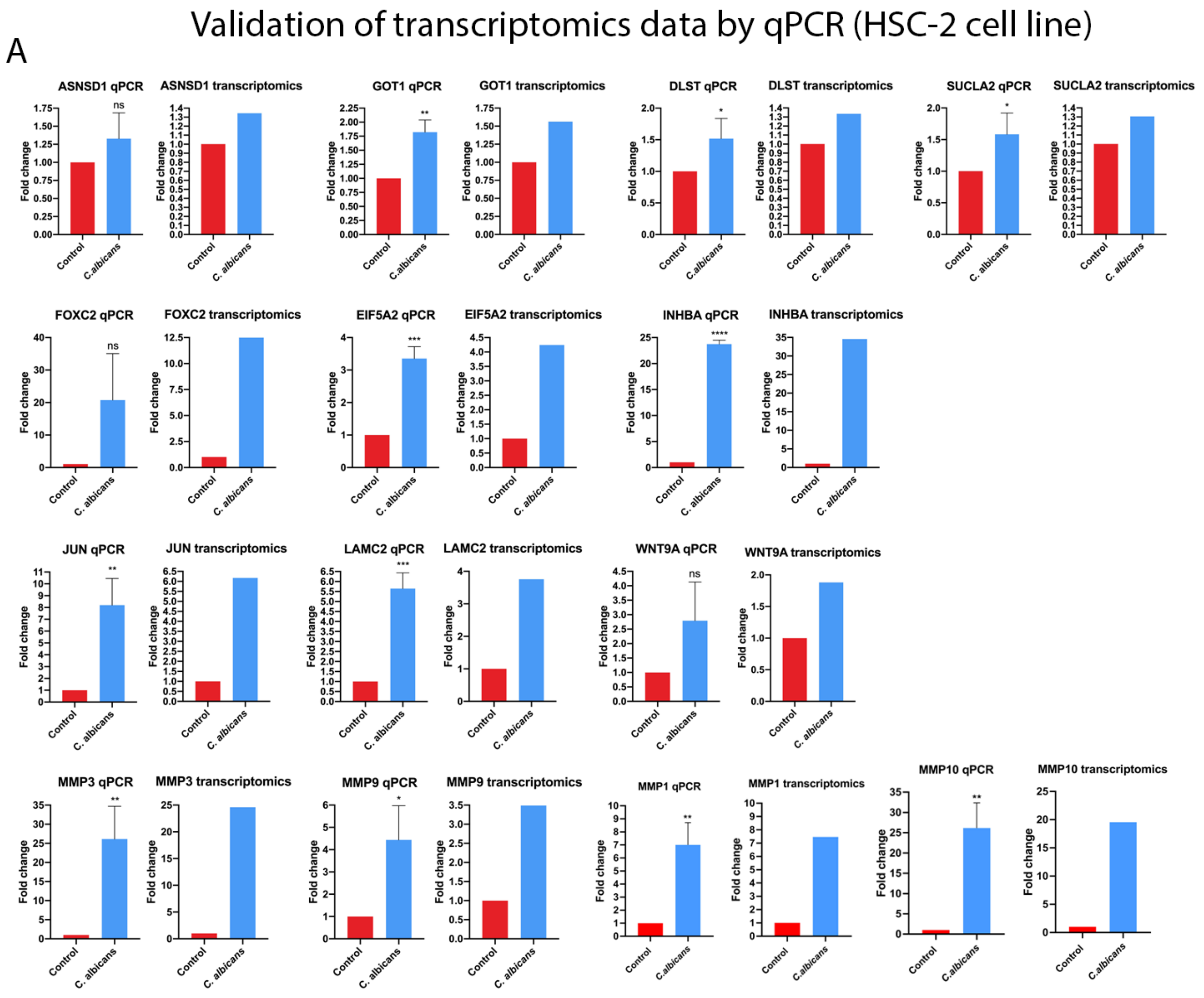


**Supp Fig9**

(A) Histopathological samples of *Candida*-colonized and *Candida*-free tumors, analysed and scored manually by a pathologist after H&E staining.

(B) Causal analyses of the genes which expression changed in OC-OSCC xenograft samples.

(C) Weight loss of the animal groups of OSCC xenograft and OC-OSCC xenograft.

(D) p63 staining of OSCC xenograft and OC-OSCC xenograft sections.

(E) E-cadherin staining of OSCC xenograft and OC-OSCC xenograft sections.

(F) Vinentin staining of OSCC xenograft and OC-OSCC xenograft sections.

A

| | OSCC Xen.1 | OC-OSCC Xen.1 | OSCC Xen.2 | OC-OSCC Xen.2 | OSCC Xen.3 | OC-OSCC Xen.3 | OSCC Xen.4 | OC-OSCC Xen.4 |
|---|---|---|---|---|---|---|---|---|
| Inflammation in surface epithelium | 0 | 1 | 0 | 3 | 0 | 2 | 0 | 2 |
| Necrosis | 0 | 1 | 1 | 1 | 1 | 1 | 1 | 1 |
| Pushing (P) or invasive (I) tumor edge | I | P/I | P | I | I | I | P | I |
| Budding / EMT | 1 | 1 | 2 | 1 | 1 | 2 | 1 | 2 |
| Invasion | 1 | 1 | 1 | 1 | 1 | 1 | 2 | 2 |
| Thrombosis | 0 | 1 | 1 | 2 | 0 | 1 | 0 | 0 |
| Peritumoral inflammation | 3g | 1l/g | 1l/g | 1l/g | 1l/g | 1l/g | 1l/g | 2l/g |

Inflammation in surface epithelium: 0 / 1 / 2 / 3
Necrosis : 0 / 1
Budding / EMT: 0 / 1 / 2
Invasion: 0 / 1 / 2 / 3
Thrombosis: 0 / 1
Peritumoral inflammation: 0 / 1 /2 /3
(g=granulocyte; l=lymphocyte)

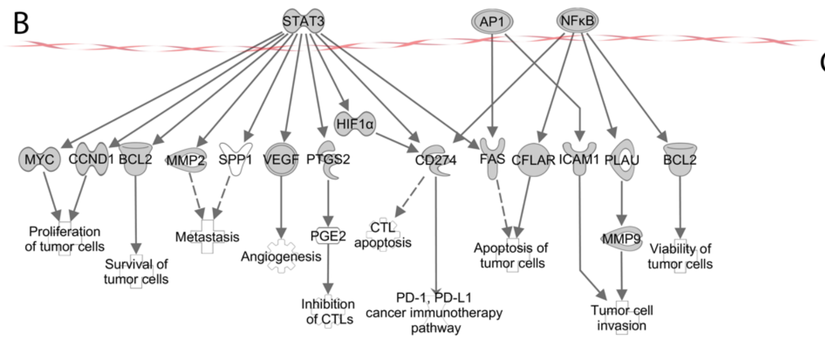


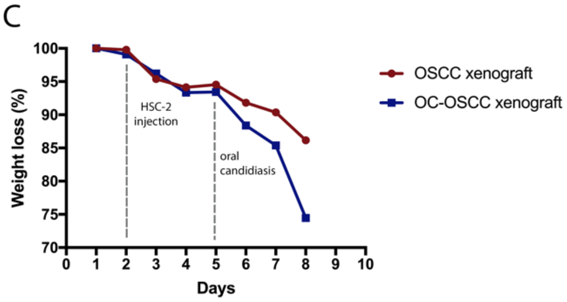


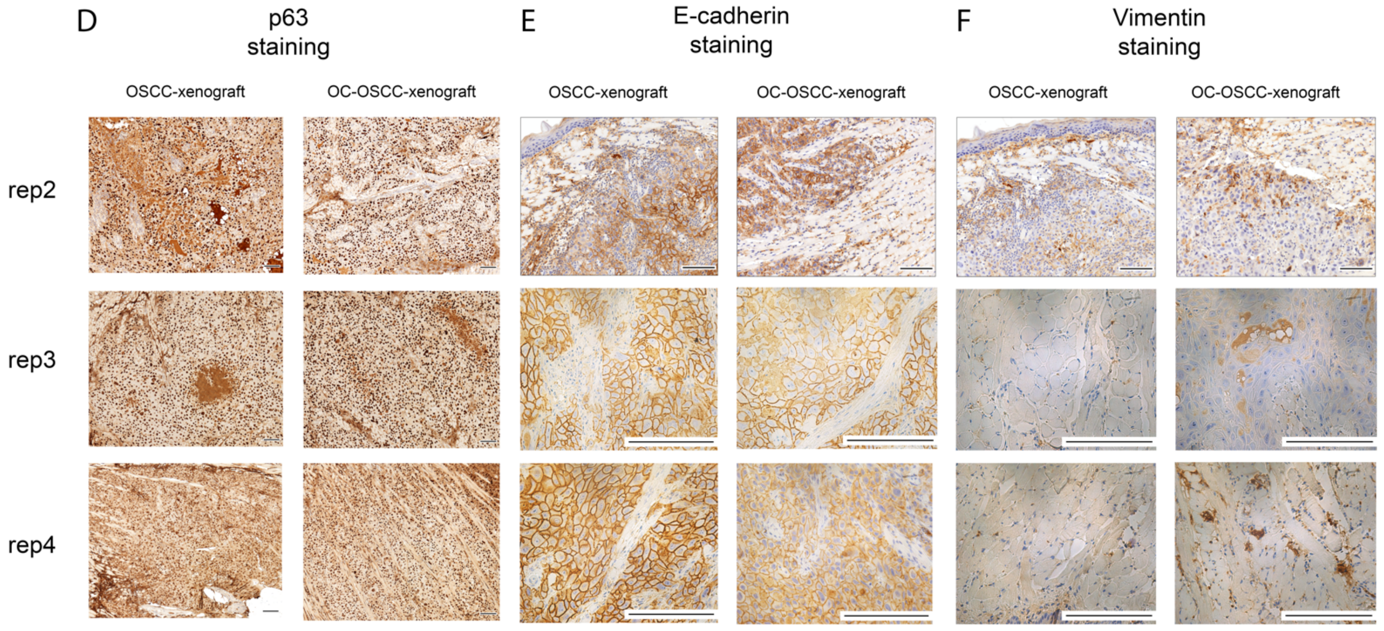


**Supp Fig10**

(A) Scoring OSCC and OC-OSCC xenograft samples manually by a pathologist.

(B) Causal analyses of the genes which expression changed in OC-OSCC samples after oral candidiasis.

(C) Weight loss of the animals after HSC-2 tumor cell injection (OSCC xenograft) and HSC-2 injection combined with oral candidiasis (OC-OSCC xenograft).

(D) p63 staining of the histopathological samples (animal 2, 3, 4)

(E) E-cadherin staining of the histopathological samples (animal 2, 3, 4)

(F) vimentin staining of the histopathological samples (animal 2, 3, 4)

A

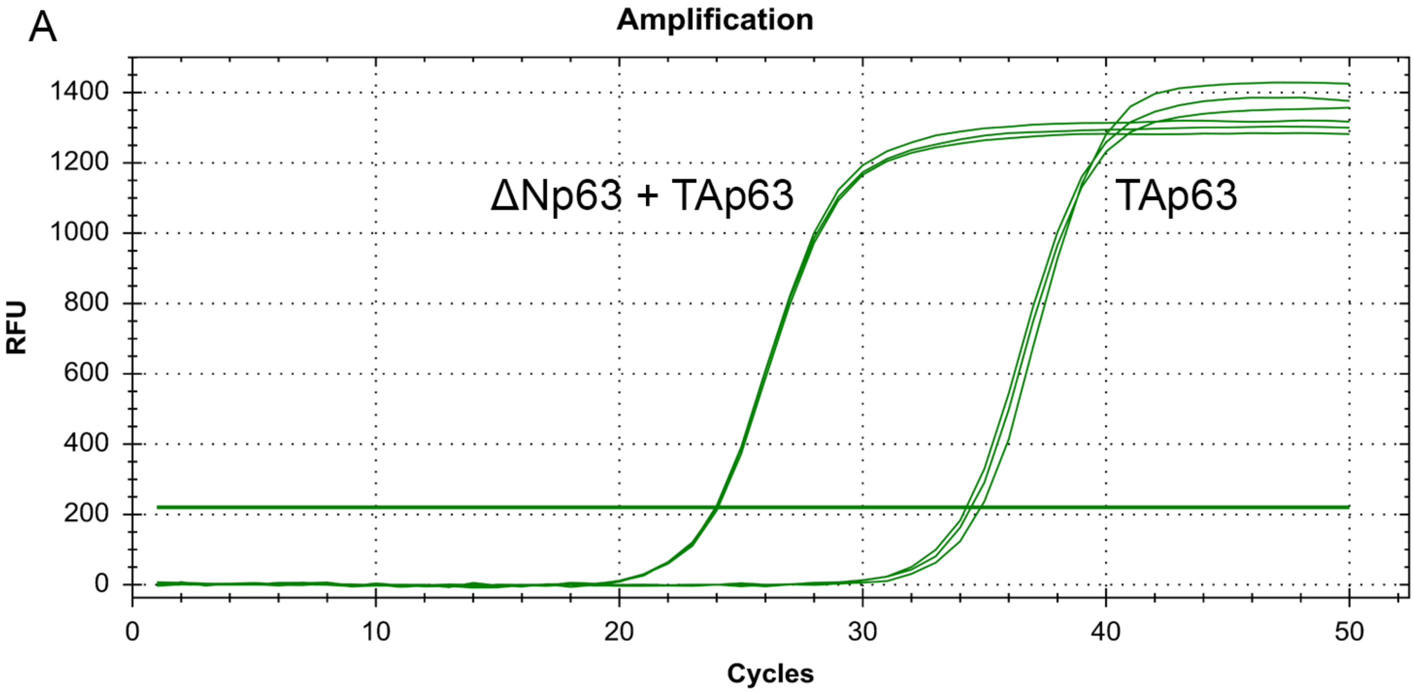


B

| | qC |
|---|---|
| ΔNp63+TAp63 | 24.02 |
| ΔNp63+TAp63 | 23.96 |
| ΔNp63+TAp63 | 24.06 |
| TAp63 | 34.45 |
| TAp63 | 34.26 |
| TAp63 | 34.85 |

**Supp Fig11**

(A) qPCR curve of TAp63 and ΔNp63+TAp63 splice variants. First primer pair was designed colse to C-terminal region, second primer pair in the N-terminal region. First primer pair amplifies both splice variant group (ΔNp63+TAp63), secon primer pair amplifies only splice variant posessing N-terminal region (TAp63).

(B) qC value of the transcript variants.

**Supp Table 1**

The precursor mass, fragment ion mass, polarity, retention time and fragmentation energy of the metabolites

**Supp Table 2**

Differentially expressed genes. *In vitro* and *in vivo* sequencing data.

**Supp Table 3**

qPCR primer list of the genes which were validated

**Supp Table 4**

120 genes derived from literature data (25 papers) which are involved in OSCC progression.
